# Vitamin D modulation of mitochondrial oxidative metabolism and mTOR enforces stress tolerance and anti-cancer responses

**DOI:** 10.1101/2021.06.08.447559

**Authors:** Mikayla Quigley, Sandra Rieger, Enrico Capobianco, Zheng Wang, Hengguang Zhao, Martin Hewison, Thomas S. Lisse

## Abstract

The relationship between vitamin D and reactive oxygen species (ROS), two integral signaling and damaging molecules of the cell, is poorly understood. This is striking, given that both factors are involved in cancer cell regulation and metabolism. Mitochondria (mt) dysfunction is one of the main drivers of cancer, producing higher cellular energy and ROS that can enhance oxidative stress and stress tolerance responses. To study the effects of vitamin D on metabolic and mt dysfunction, we used the vitamin D receptor (VDR)-sensitive MG-63 osteosarcoma cell model. Using biochemical approaches, active vitamin D (1,25-dihydroxyvitamin D, 1,25(OH)_2_D_3_) decreased mt ROS levels, membrane potential (ΔΨ_mt_), biogenesis, and translation, while enforcing endoplasmic reticulum/mitohormetic stress tolerance responses. Using a mitochondria-focused transcriptomic approach, gene set enrichment and pathway analyses show that 1,25(OH)_2_D_3_ lowered mt fusion/fission and oxidative phosphorylation (OXPHOS). By contrast, mitophagy, ROS defense, and epigenetic gene regulation were enhanced after 1,25(OH)_2_D_3_ treatment, as well as key metabolic enzymes that regulate fluxes of substrates for cellular architecture and a shift toward non-oxidative energy metabolism. ATACseq revealed putative oxi-sensitive and tumor-suppressing transcription factors that may regulate important mt functional genes such as the mTORC1 inhibitor, *DDIT4/REDD1*. DDIT4/REDD1 was predominantly localized to the outer mt membrane in untreated MG-63 cells yet sequestered in the cytoplasm after 1,25(OH)_2_D_3_ and rotenone treatments, suggesting a level of control by membrane depolarization to facilitate its cytoplasmic mTORC1 inhibitory function. The results show that vitamin D activates distinct adaptive metabolic responses involving mitochondria to regain redox balance and control the growth of cancer cells.

## Introduction

The altered metabolism of cancer cells imparts a large impact on redox (reduction-oxidation) homeostasis resulting in enhanced production of reactive oxygen species (ROS) that drives and sustains cancer cell proliferation and evasion^1^. One counter measurement to elevated ROS is the increase of antioxidants that allows cancer cells to persist in a damaging environment.

However, as cancer cells persist in the continuing pro-oxidant environment, DNA is further damaged and the genome becomes more unstable with concomitant metabolic reprogramming^1^. Over time, the metabolic reprogramming further distances cancer cells from redox homeostasis, leading to a vicious cycle of accumulating genetic lesions toward cancer progression. Understanding the mechanisms and how to control the dysfunctional and dysregulated metabolism of cancer cells is a major challenge in cancer biology.

Perturbation of the vitamin D metabolic and signaling systems in humans and animals are associated with several diseases and disorders including alopeicia^2, 3^, osteosarcopenia^4^, diabetes^5^, and cancer^6^. Vitamin D cellular activities are regulated primarily by the hydroxylated metabolite 1-alpha, 25-dihydroxyvitamin D_3_ (also called 1,25(OH)_2_D_3_ or calcitriol). 1,25(OH)_2_D_3_ effects are mediated by the vitamin D receptor (VDR), a member of the intracellular nuclear receptor superfamily^7, 8^. A large body of studies has all implicated a suppressive role of vitamin D in cancer development and improved cancer patient and animal survival^6, 9–12^. For example, elevated serum vitamin D levels at diagnosis have been linked to extended survival rates in cancer patients^13^. And most recently, a large clinical trial involving vitamin D supplementation suggests that vitamin D can benefit patients with advanced or lethal cancers by prolonging mortality and survival in the study population^10, 14^. Despite these encouraging results from the clinical trial, the precise mechanism for anti-cancer effects of vitamin D remains elusive.

Understanding how what we eat (e.g., fruits and vegetables, which are rich in antioxidants like vitamin C^15^), what we expose ourselves to (e.g., sunlight, pollution), and the metabolism that can lead to energy-reducing equivalents that drive cellular replication and differentiation at the genetic level provides insight toward the circle of life. In contrast, understanding how molecular factors such as ROS and their antioxidants can regulate the cell cycle, differentiation, and adaptive responses (e.g., DNA damage, mutagenesis) at the genetic level provides insight toward the circle of death. Numerous mechanisms exist to help dictate the consequences of cellular oxidative stress. For example, the cytoprotective upregulation of DNA repair transcripts allow for DNA mismatch repair, non-homologous end-joining, and base excision repair in cells to handle the high levels of oxidative damage. Other avenues may consist of oxidation of membrane receptors, signaling molecules, redox-sensitive transcription factors, and epigenetic transcriptional regulators including histone deacetylase family members to regulate the cell’s response to stress and cancer development^16^. Understanding the metabolic oxidation/reduction reactions and cellular responses to various biological and environmental factors remains a major milestone in cancer biology and therapy.

Thus, in the present study, we postulated that effects on the metabolic oxidation/reduction reactions that are characteristic of cancer cells may be crucial to the anti-cancer effects of the environmental/nutritional factor vitamin D. We applied genome-wide transcriptomic and epigenomic approaches to provide a comprehensive understanding of how vitamin D modulates mitochondrial functions and counteracts tumorigenicity within the MG-63 osteosarcoma cell model, followed by mitochondrial biochemical and ultrastructural investigations. Based on these approaches, we provide evidence of novel regulators of mitochondrial stress, biogenesis, translation, organellar hormesis, and cancer metabolic fates mediated by vitamin D, and cooperating factors leading to an overall reduction in oxidative stress levels. In particular, although the tumor suppressor DNA damage inducible transcript 4 (DDIT4) is induced under stress conditions in normal bone cells to inhibit metabolism activated by the mammalian target of rapamycin (mTOR) in the cytoplasm^17^, MG-63 osteosarcoma cells exhibit high levels of DDIT4 sequestered to the mitochondria as a potential mechanism to regulate mTOR activation and cancer progression. In contrast, vitamin D treatment of MG-63 cells increased the expression and cytoplasmic localization of DDIT4 through separation from the outer mitochondrial membrane. Ironically, a meta-analysis of numerous cancer cell types identified DDIT4 as being overexpressed compared to non-cancerous cells and associated with poor survival outcomes despite being a potent mTOR inhibitor^18^. Based on our findings from MG-63 cells, we propose that vitamin D may suppress tumor progression of other cancer types that involves mitochondrial-to-cytoplasmic DDIT4/REDD1 exchange. Overall, the results herein establish that vitamin D can target the deregulation of specific metabolic hubs within osteosarcoma cells to suppress tumorigenicity, thus attempting to maintain the delicate balance between the cycle of life and death.

## Results

### Genome-wide assessment of vitamin D-mediated transcription using RNAseq

Previous studies have shown that 1,25(OH)_2_D_3_ (called vitamin D from hereon) can suppress the growth of MG-63 cells, but not of receptor-poor cell lines in standard 2D culture assays within the range of 100nM (10^-8^M or 40ng/mL) and 10nM (10^-9^M or 4ng/mL)^19^. However, 3D colony formation in soft agar is the gold-standard assay to study anchorage-independent carcinogenesis *in vitro*^20^, and has not been tested on vitamin D treated MG-63 cells to date. After 14 days of continuous vitamin D treatment, both 10nM and 100nM of vitamin D resulted in a significant reduction in overall colony size (**Figure 1A,B**). However, 10nM of vitamin D did not significantly affect the number of colonies after treatment, in contrast to 100nM (**Figure 1B**). This suggests that vitamin D may induce cell apoptosis at the higher 100nM concentration, and that vitamin D can suppress MG-63 tumor growth at 10nM via non-apoptotic mechanisms and lower toxicity (see later).

**Figure 1.**
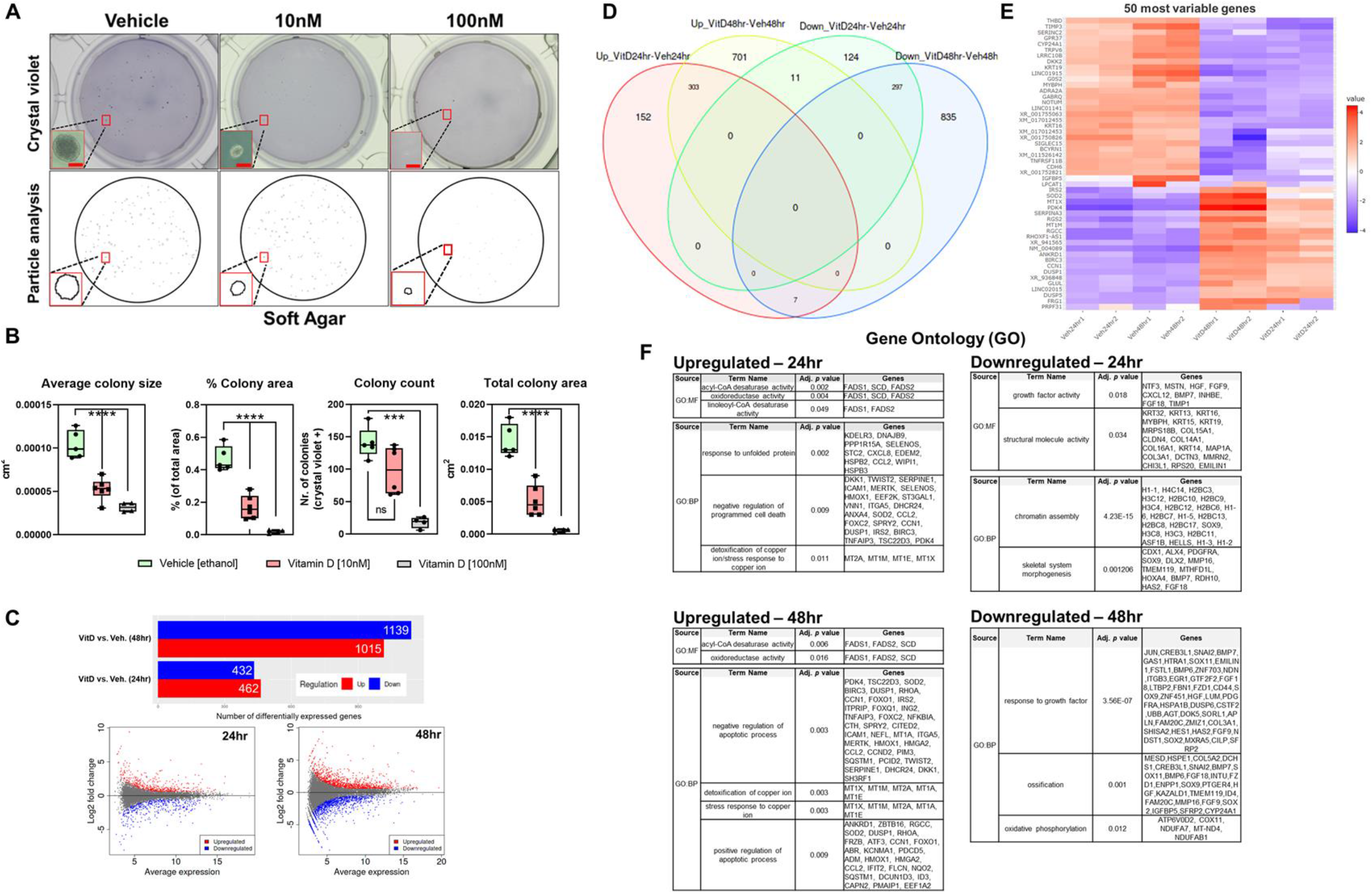
**Genome-wide assessment of vitamin D-mediated transcription using RNAseq** A) Top: Representative macroscopic images of soft agar colony formation of MG-63 cells treated with 1,25(OH)_2_D_3_ for 14 days. Bar=100µm. Bottom: ImageJ particle analysis of colonies. B) Quantitation of the data from A, summed from five-six representative macroscopic fields for each condition using data derived from ImageJ (n=5-6). Data are presented as mean±SEM error bars; ^∗∗∗∗^*p* ≤ 0.0001, and ^∗∗∗^*p* ≤ 0.001 (one-way ANOVA with Tukey’s multiple comparisons test). C) MA plot and summary of differentially expressed genes (DEGs) based on DESeq2 method of RNAseq data. Plotted are the differences between measurements from vitamin D [10nM] versus vehicle-treated cells by transforming the data onto M (log ratio) and A (mean average) scales. DEGs selected based on false discovery rate (FDR) set to 0.05 and logFoldChange(1) compared to controls. D) Venn analysis of overlapping gene sets performed at http://www.interactivenn.net. E) Heatmap with hierarchical clustering tree of the 50 most variable genes (n=2 samples/condition). F) Functional annotation and enrichment analysis using Gene Ontology (GO). Annotated genes, descriptors, and adjusted *p* values (≤0.05 considered significant) presented. GO molecular function (MF) and biological process (BP) domains are used to establish relationships. DEGs filtered using g:GOSt at https://biit.cs.ut.ee/gprofiler/gost.

To define the genome-wide impact on transcription by vitamin D on MG-63 cells under the growth-inhibiting conditions at 10nM, high throughput RNAseq analysis was carried out. After 24 hours of continuous vitamin D treatment, there were 462 and 432 differentially up and downregulated genes, respectively, compared to vehicle treatment using a false discovery rate (FDR) threshold of 0.05 and log Fold Change(1) with a high degree of correlation (Pearson correlation coefficient = 0.96-0.99) (**Figure 1C, S1, & Worksheet S1**). After 48 hours of vitamin D treatment, there were 1,015 and 1,139 differentially up and downregulated genes, respectively. Venn analysis revealed that there were 152 and 124 uniquely up and downregulated genes, respectively, mediated by vitamin D after 24 hours (**Figure 1D**). While 701 and 835 genes were uniquely up and downregulated, respectively, relative to 48 hours of vitamin D treatment. To identify subgroups of genes that share expression patterns, we ranked genes by their standard deviation to obtain hierarchical clusters using DESeq2 (**Figure 1E, Worksheet S2**). An interactive heatmap of the 50-most variable genes shows that vitamin D induced genes such as *SOD2*, *IRS2*, *BIRC3*, and *DUSP1/5*, which are either cytoplasmic or mitochondrial signaling molecules that mediate the effects of growth factors and/or cytokine interactions with known anti-cancer properties^21^. For example, vitamin D strongly induced the expression of the mitochondrial, but not cytosolic, manganese superoxide dismutase, SOD2, which converts the free radical O_2_^• −^ (superoxide) to H_2_O_2_ to defend against free radicals. The 50-most variable downregulated genes included the cytochrome P450 family 24 subfamily A member 1 (*CYP24A1*), the onco-channel *TRPV6*, and *DKK2* (i.e., a Wnt mediator of tumor immune evasion). CYP24A1 functions as a mitochondrial monooxygenase that catalyzes the 24-hydroxylation and catabolism of vitamin D, suggesting a negative feedback response to preserve vitamin D signaling and its anti-cancer effects in MG63 cells.

Next, we studied the relationships among the up and down-regulated genes relative to their enriched Gene Ontology (GO) terms using hierarchical clustering trees (**Figure S2A**). The downregulated genes were overwhelmingly involved in *nucleosome/chromatin assembly* and *organization*, as well as *DNA replication*. Since nucleosomes assemble and become octameric during DNA replication amassing on daughter DNA strands, these findings suggest vitamin decreases replication, replication stress, and genomic instability associated with cancer. The decrease in nucleosome assembly also suggests that vitamin D-treated MG-63 cells may be in interphase of the cell cycle, supporting our previous studies^17^, where DNA is less compact and associated with increased chromatin accessibility (see later). Although *chromatin remodeling*, per se, was not a feature of the analysis, we did identify chromatin-modifying enzymes (e.g., SIRT1,4) that were differentially regulated after vitamin D treatment. Nevertheless, the upregulated genes were associated with *programmed cell death*, *translation*, and *response to organic substance*. Of note, although regulators of apoptotic pathways were found to be enriched, we observed no changes in the early apoptosis marker Annexin V phosphatidylserine in vitamin D-treated MG-63 cells at 10nM (data not shown). We also used the dimension reduction algorithm, t-SNE, to map the top genes, and then identified four clusters of enriched pathways called k-means that were further mapped to GO biological processes (**Figure S2B, Worksheet S3**). Cluster A consisted of genes upregulated after 48-hour of vitamin D treatment that was enriched for the *defense response to virus* pathway. Cluster B consisted of genes upregulated after vitamin D treatment for both 24 and 48 hours that were enriched for the *stress response* pathway. Cluster C consisted of genes downregulated after 48-hour vitamin D treatment that enriched for the *chromosome organization* pathway. Lastly, Cluster D consisted of genes downregulated after both 24 and 48 hours that were enriched for *chromatin/nucleosome assembly* and *cell development* pathways. These findings show that vitamin D regulates genome architecture and downstream stress response pathways as part of its anti-cancer response.

### Functional enrichment analysis reveals vitamin D-mediated cancer inhibition via mitochondrial OXPHOS and stress regulators

Functional annotation and gene set enrichment analysis (GSEA) were performed using several methods to reflect the heterogeneity of data repositories and statistical approaches. We first used the g:GOSt program to map genes to known functional information to determine statistically significant enriched relationships. The data were stratified based on GO molecular functions (MF), biological processes (BP), and cellular components (**Worksheet S4, S5**). Based on GO-MF subset analysis, genes that regulate fatty acid desaturases were upregulated after vitamin D treatment, suggesting a putative role in unsaturated fatty acid biosynthesis and utilization (**Figure 1F**). Based on GO-BP, vitamin D treatment induced genes that regulate *unfolded proteins*, *programmed cell death*, and the *detoxification of metal ions*. On the other hand, vitamin D suppressed *growth factors and structural molecule activity-related* genes based on GO-MF. Based on GO-BP, vitamin D suppressed *chromatin assembly*, *morphogenesis,* and *oxidative phosphorylation* (OXPHOS)-related genes. The OXPHOS genes include *COX11*, which is a copper-binding subunit of the cytochrome c oxidase enzyme in the electron transfer chain in the mitochondria. Several respiratory chain NADH dehydrogenase subunit genes were also downregulated, including *NDUFA7*, *MT-ND4*, and *NDUFAB1*. The remainder of the enriched pathways with large gene sets are included in **Worksheets S4, S5**, which also include genes enriched for *telomere maintenance* and *adipogenesis*, for example.

We next performed GSEA in conjunction with the Molecular Signatures Database (MSigDB, version 7.3) of annotated gene sets. GSEA associates a treatment phenotype to a group or a list of weighted genes for comparison. The MSigDB gene sets are divided into nine major collections, whereby the Hallmarks (H) gene sets (i.e., 50 gene sets) and Canonical Pathways (CP) gene sets (i.e., 189 gene sets) were applied with the cut-off of *p*-value ≤ 0.05 to select biologically meaningful processes (**Worksheet S6**). GSEA analysis of the 24-hour vitamin D-treated samples revealed gene sets related to *inflammation*, *hypoxia*, and *epithelial-mesenchymal transition* (EMT) pathways that were not discovered using g:GOSt (**Figure 2A**). For example, the data suggests that vitamin D can reverse EMT to suppress mesenchymal metastasis through downregulation of SNAI2, a key zinc finger transcription factor that maintains the loose mesenchymal phenotype (**Figure 2A,B**). After 48 hours of vitamin D treatment, the enriched pathways were related to *hypoxia*, *glycolysis*, *inflammation*, *<u>unfolded protein response</u>*<u>, *mTOR pathway*, c*holesterol homeostasis*, *apoptosis*, *xenobiotic metabolism,*</u> and *p53 signaling* (**Figure 2B**). Key upregulated genes include *<u>DDIT4/REDD1</u>* and sequestosome 1 (SQSTM1), which target the direct inhibition of mTOR or indirect effects through autophagy, respectively. In terms of hypoxia, decreased OXPHOS after vitamin D treatment is likely to increase molecular oxygen levels as hypoxia in cancer cells is partly due to increasing O_2_ consumption and reduction to water that can thereby induce EMT^22^. Hyperoxia is also supported by the increased SOD2 levels after vitamin D treatment, as SOD2 metabolizes superoxide radicals into hydrogen peroxide. These findings suggest that vitamin D affects major pathways involved in oxygen levels and the growth regulation of tumor cells.

**Figure 2.**
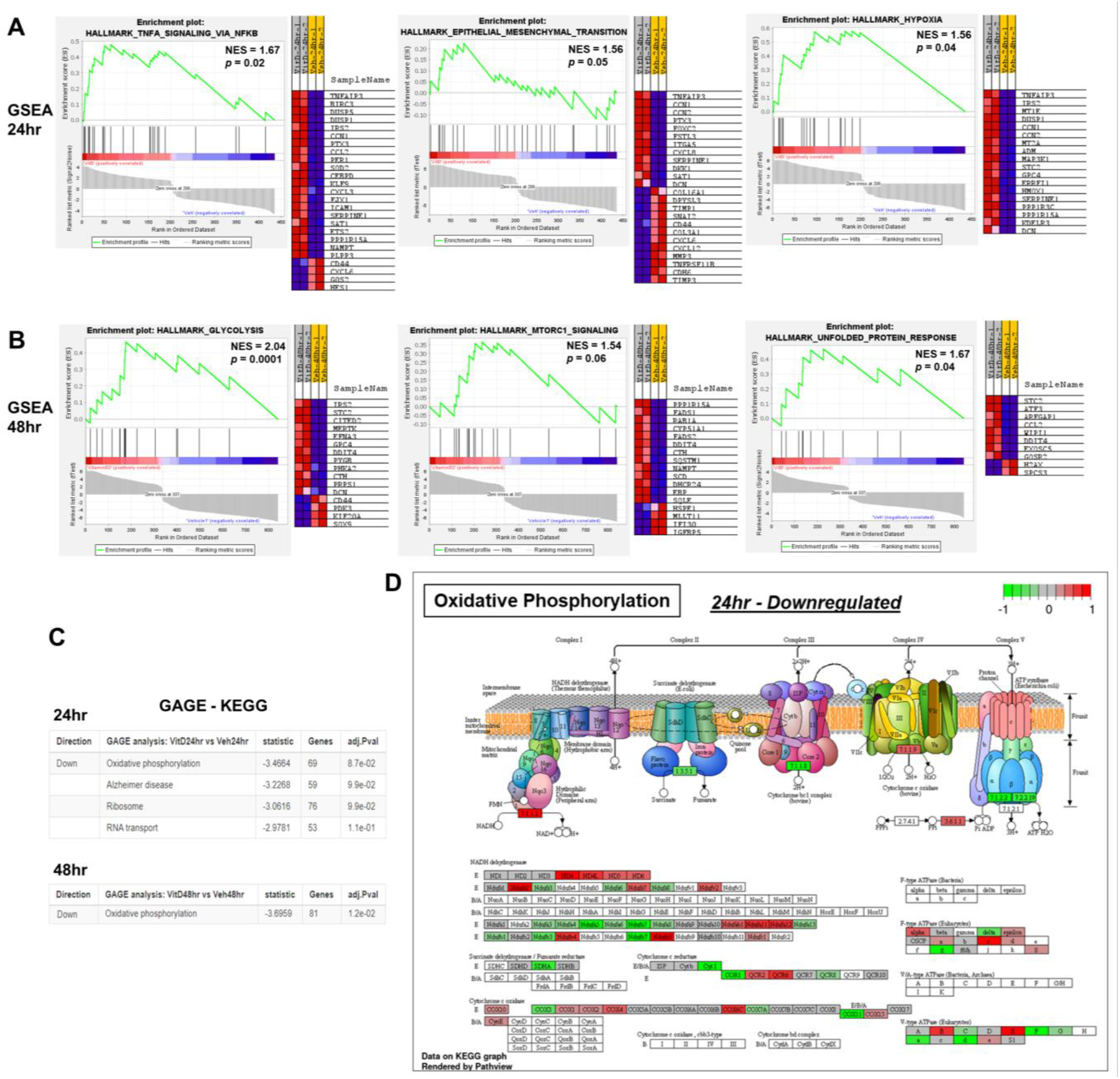
**GSEA analysis of vitamin D treated MG-63 cells** A) Gene Set Enrichment Analysis (GSEA) to identify pathways enriched in ranked gene lists after 24 hours of vitamin D treatment. GSEA score curves depict the strength of gene sets in the Molecular Signature Database. “Signal-to-Noise” ratio (SNR) statistic was used to rank the genes per their correlation with either vitamin D [10nM] treatment (red) or vehicle treatment (blue). Generally, the gene sets will be significant when a proportionally large number of genes fall in the upper or lower part of the distribution. The heatmap on the right of each panel depicts the genes contributing to the enriched pathway. The green curve corresponds to the enrichment score, which is the running sum of the weighted enrichment score obtained from GSEA. Positive normalized enrichment score (NES) and significant *p* values denote the most enriched pathways of the members of the gene set. Full pathway and gene list in Worksheet S7, GSEA performed at http://www.broad.mit.edu/gsea. B) GAGE method for gene set enrichment of vitamin D treatment. The fold change (log-based) was used for the *per* gene statistics and adjusted *p* values (≤0.05 considered significant) presented. Full pathway and gene list in Worksheet S8, GAGE performed at https://pathview.uncc.edu/gageIndex.

In addition, we applied Generally Applicable Gene-set Enrichment (GAGE) analysis that has no limitations on sample size based on a parametric gene randomization method to test the significance of gene sets using log-based fold changes as the *per gene* statistic. By using the absolute values of fold change in the GAGE analysis combined with the Kyoto Encyclopedia of Genes and Genomes (KEGG) database, vitamin D was shown to significantly downregulate a more dynamic OXPHOS gene set at both 24 and 48 hours of treatment (**Figure 2C and Worksheet S7**). The GAGE output was shared with Pathview to rationalize the OXPHOS genes (**Figure 2D**), whereby the analysis shows that vitamin D downregulates the second (II) and third (III) large enzyme complexes (i.e., succinate dehydrogenase and cytochrome bc1 complex, respectively) in the respiratory electron transport chain of MG-63 cells. The cytochrome bc1 complex is responsible for the proton gradient as well as for the formation of O_2_^.-^. Disruption of the flow of electrons across the membrane is predicted to alter the transmembrane difference of proton electrochemical potential which ATP synthases use to generate energy (see later). Furthermore, vitamin D downregulated ATP synthase components (e.g., ATP5D, ATP5A1, ATP5C1) as another mode to regulate overall mitochondrial activity. Collectively, these findings suggest that vitamin D promotes additional metabolic shifts that involve the suppression of mitochondrial OXPHOS as part of its anti-cancer strategy.

### Vitamin D-mediated organellar hormesis enforces stress tolerance and growth inhibition of MG-63 cells

Our genome-wide bioinformatics analysis suggests the involvement of the unfolded protein response (UPR) in vitamin D-treated MG-63 cells, which is known to mediate stress tolerance and organismal longevity involving a process called hormesis^23^. Hormesis describes a phenomenon where mild cellular stress caused by unfolded proteins stimulates alternative signaling pathways with beneficial, over-compensating outcomes and organellar connectivity^23^. To better understand how vitamin D modulates hormetic responses in cancer cells, we investigated various ER and mitochondrial hormetic signaling pathways that involve antioxidants and protein-folding chaperones (**Figure 3**). The ER transmembrane receptor protein kinases (ER-TRK) IRE1 and PERK, and the transcription factor ATF6 govern the expression of factors that protect cells as part of the hormetic UPR by promoting cell cycle arrest, protein translation inhibition, and chaperone production. A proxy for activated IRE1 is cleavage of 26 base pairs from its substrate, XBP1, to generate a spliced form called sXBP1 that functions as a transcription factor for expression of binding immunoglobulin protein (BIP, also called GRP78 or HSPA5), which functions as a major ER stress chaperone. To characterize UPR in the MG-63 cell system, thapsigargin and tunicamycin (i.e., blockers of the ER ATPase/SERCA pump and glycoprotein synthesis, respectively) were first used and found to induce a dose-dependent increase in sXBP1 and BIP/HSPA5 (**Figure 3A-C**). Interestingly, vitamin D treatment enhanced sXBP1 in a time-dependent manner at 10nM, but not at 100nM (**Figure 3D,E**) with no change in BIP mRNA levels across all concentrations suggesting a hormetic response to insoluble proteins (**Figure 3G,H**). As the proxies for ATF6 activation are upregulation of BIP and uXBP1, our findings also suggest that ATF6 plays a minimal role in the vitamin D response (**Figure 3E**). Two proxies for PERK activation are ATF4 and CHOP (also called DDIT3 or GADD153), whereby RNAseq analysis showed no changes in both transcripts after vitamin D treatment (**Figure 3H** and **Worksheet S1**). Therefore, we investigated potential hormetic antioxidative responses of the alternative ER-TRK, recently described in *C. elegans*^24^, in the context of vitamin D by appraising the human glutathione S-transferase family of genes. We only observed statistically significant increases in glutathione S-transferase kappa 1 (GSTK1) and glutathione S-transferase Mu 4 (GSTM4) after vitamin D treatment of MG-63 cells (**Figure 3I**), whereby lower levels of GSTK1 have been linked to the elevation of mt ROS underlying hypertrophic cardiomyopathy^25^. Lastly, since the bioinformatics analysis also suggests the downregulation of OXPHOS, we assessed mitochondrial UPR by way of activating transcription factor 5 (ATF5) (**Figure 3J**). ATF5 is a major mitochondrial stress regulator that can induce proteostasis and chaperonin production^26^, whereby 10nM of vitamin D treatment significantly downregulated ATF5 in MG-63 cells, the effect of which dissipated at higher concentrations signifying a hormetic response (**Figure 3J**). Overall, the results suggest that vitamin D activates distinct hormetic adaptive responses in the ER and mitochondria to regain control of the growth of cancer cells, which may underly beneficial interorganellar communication to overcome cancer stress (**Figure 3K**).

**Figure 3.**
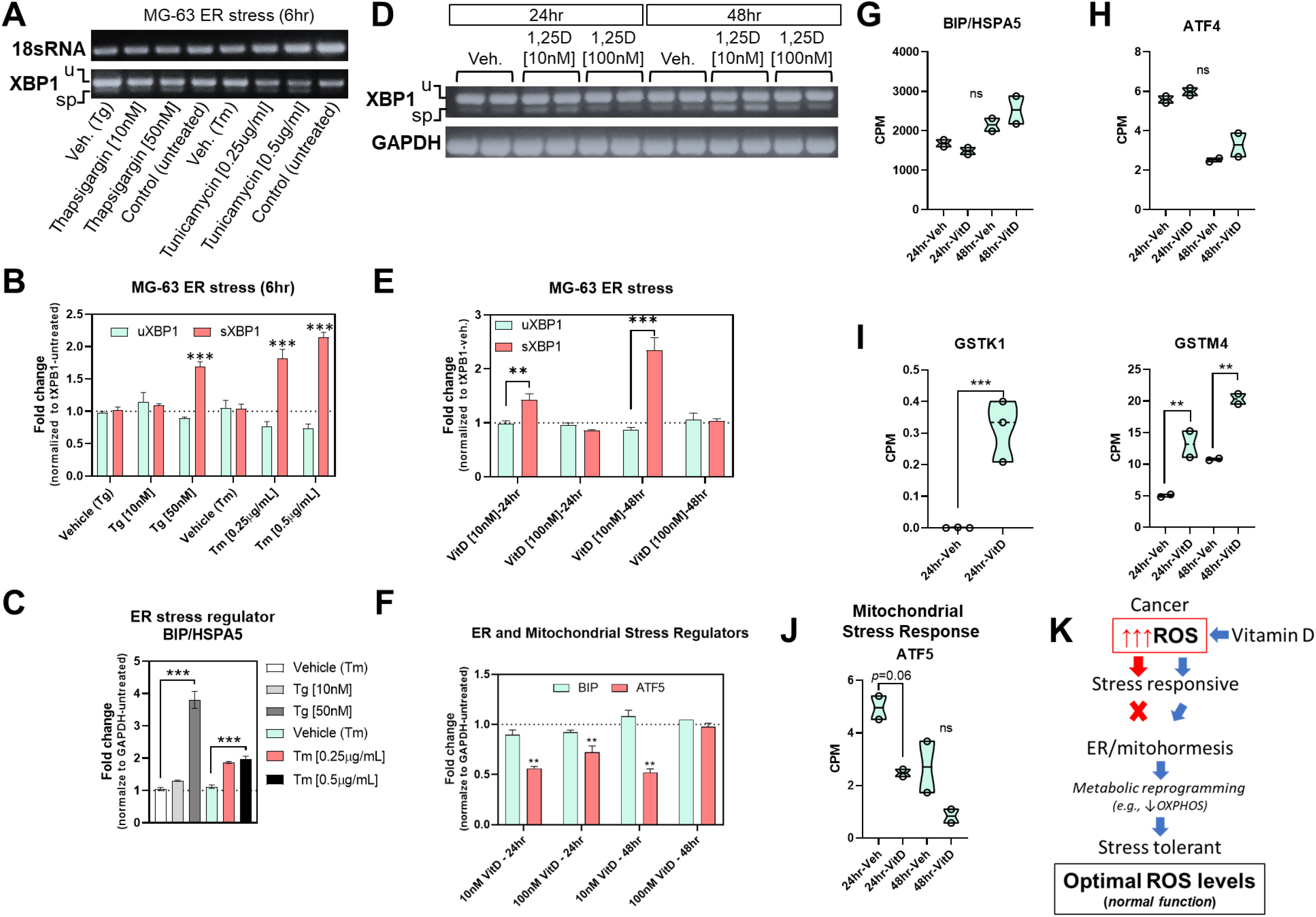
**Vitamin D and ER/Mitochondrial stress regulation** A) Representative end-point PCR analysis of IRE1-XBP1 expression after 6 hours of positive control treatments. u (unspliced XBP1; 256bp), sp (spliced XBP1; 230bp), 18sRNA (190bp). B) Real-time PCR analysis of IRE1-XBP1 expression after 6 hours of positive control treatments. The graph depicts fold change of either uXBP1 (unspliced) or sXBP1 (spliced) normalized to the total XBP1 levels. Data are presented as mean±SEM error bars (n=3 samples/condition); ^∗∗∗^*p* ≤ 0.001 (one-way ANOVA with Tukey’s multiple comparisons test compared to respective vehicle). C) Real-time PCR analysis of BIP/HSPA5 expression in positive controls. Data are presented as mean±SEM error bars (n=3 samples/condition); ^∗∗∗^*p* ≤ 0.001 (one-way ANOVA with Tukey’s multiple comparisons test compared to respective vehicle). D) Representative end-point PCR analysis of IRE1-XBP1 expression after 24-48 hours of vitamin D treatments. u (unspliced XBP1; 256bp), sp (spliced XBP1; 230bp), GAPDH (350bp). E) Real-time PCR analysis of IRE1-XBP1 expression after 24-48 hours of vitamin D treatments. The graph depicts fold change of either uXBP1 (unspliced) or sXBP1 (spliced) normalized to the total XBP1 levels. Data are presented as mean±SEM error bars (n=3 samples/condition); ^∗∗∗^*p* ≤ 0.001, ^∗∗^*p* ≤ 0.01 (one-way ANOVA with Tukey’s multiple comparisons test compared to respective vehicle). F) Real-time PCR analysis of BIP/HSPA5 and ATF5 expression after 24-48 hours of vitamin D treatments. Data are presented as mean±SEM error bars (n=3 samples/condition); ^∗∗^*p* ≤ 0.01 (one-way ANOVA with Tukey’s multiple comparisons test compared to respective vehicle). G-J) RNAseq analysis of ER/mitochondrial stress and hormetic regulators. A two-way ANOVA was performed with Bonferroni’s multiple comparisons test using the counts per million (CPM) values (n=2 samples/condition), where the *p*-value summaries were depicted as ^∗∗∗∗^*p* ≤ 0.0001, ^∗∗∗^*p* ≤ 0.001, and ^∗∗^*p* ≤ 0.01. ns (not significant). K) Proposed model: Vitamin D enforces stress tolerance in cancer cells via metabolic reprogramming involving ER/mitohormesis.

### A multi-omics approach to study mitochondrial anti-cancer responses to vitamin D

Given that vitamin D suppresses mitochondrial UPR, we performed a more granular multi-omics assessment of mitochondrial transcriptional changes using the annotated databases MitoCarta and mitoXplorer. MitoCarta currently annotates 1,136 genes encoding mitochondrial proteins, while mitoXplorer contains 1,229 genes. First, we used MitoCarta (version 3.0) to identify differentially regulated mitochondria-related genes from our RNAseq data set^27^. Among the 1,477 upregulated vitamin D-mediated differentially expressed genes (DEGs) (**Figure 1**), we identified 79 genes that encode mitochondria proteins within the combined 24- and 48-hour gene sets (∼5%; **Figure 4A**, **Worksheet S8**). Among the 1,571 downregulated vitamin D-mediated DEGs (**Figure 1**), we identified 45 genes encoding mitochondrial proteins in total (∼2.8%; **Figure 4A, Worksheet S8**). However, MitoCarta provides no annotation on the genes, and to understand the biological significance behind these changes, we utilized the annotated mitoXplorer (version 1.0) necessary for pathway analysis. In all, there were 64 and 37 vitamin D-mediated up and downregulated mitochondrial genes, respectively, that were common between the two repositories (**Figure 4B**). There were only 15 and 8 up and downregulated vitamin D-mediated mitochondrial genes, respectively, that were specific to the MitoCarta repository and not included in the mitoXplorer annotative analysis. Based on the mitoXplorer analysis, the vitamin D-mediated downregulated DEGs after 24 hours included *MRPS18B*, which encodes a 28S subunit mitoribosomal protein involved in protein translation (**Figure 4C, Worksheet S8**). In addition, HSPA1A and B, members of the heat shock protein family A, were also downregulated by vitamin D, suggesting a lowering of stress aggregation and increased protein stability in mitochondria. In terms of metabolism, dimethylglycine dehydrogenase (DMGDH), a mitochondrial enzyme involved in phosphatidylcholine and lipid metabolism, and glycine modifications, was elevated after vitamin D treatment (**Figure 1F**). Recently, studies have shown that DMGDH can play a role in antioxidant defense^28^, and low levels can be a diagnostic and prognostic marker for hepatocellular carcinoma metastasis by acting on the Akt pathway^29^. Genes that regulated beta oxidation of fatty acids were also found to be suppressed by vitamin D treatment (↓ACAA2), suggesting another mean for ROS reduction^30^. Interestingly, mitochondrial amino acid metabolism and detoxification were upregulated after vitamin D treatment by way of glutamate-ammonia ligase (GLUL), which is a mitochondrial enzyme that catalyzes the synthesis of glutamine from the more toxic glutamate and ammonia. In addition, nitrilase omega-amidase (NIT2) was upregulated by vitamin D, which is known to play a role in arresting cells to remove toxic intermediates such as 2-oxoglutaramate^31^. Pyruvate metabolism was also affected after vitamin D treatment by way of upregulation of the mitochondrial pyruvate dehydrogenase kinase 4 (PDK4). PDK4 inhibits the mitochondrial pyruvate dehydrogenase complex to reduce pyruvate conversion from glucose, suggesting that vitamin D may conserve glucose metabolism (i.e., slowing glycolysis), as during hibernation, by decreasing its conversion to acetyl-CoA.

**Figure 4.**
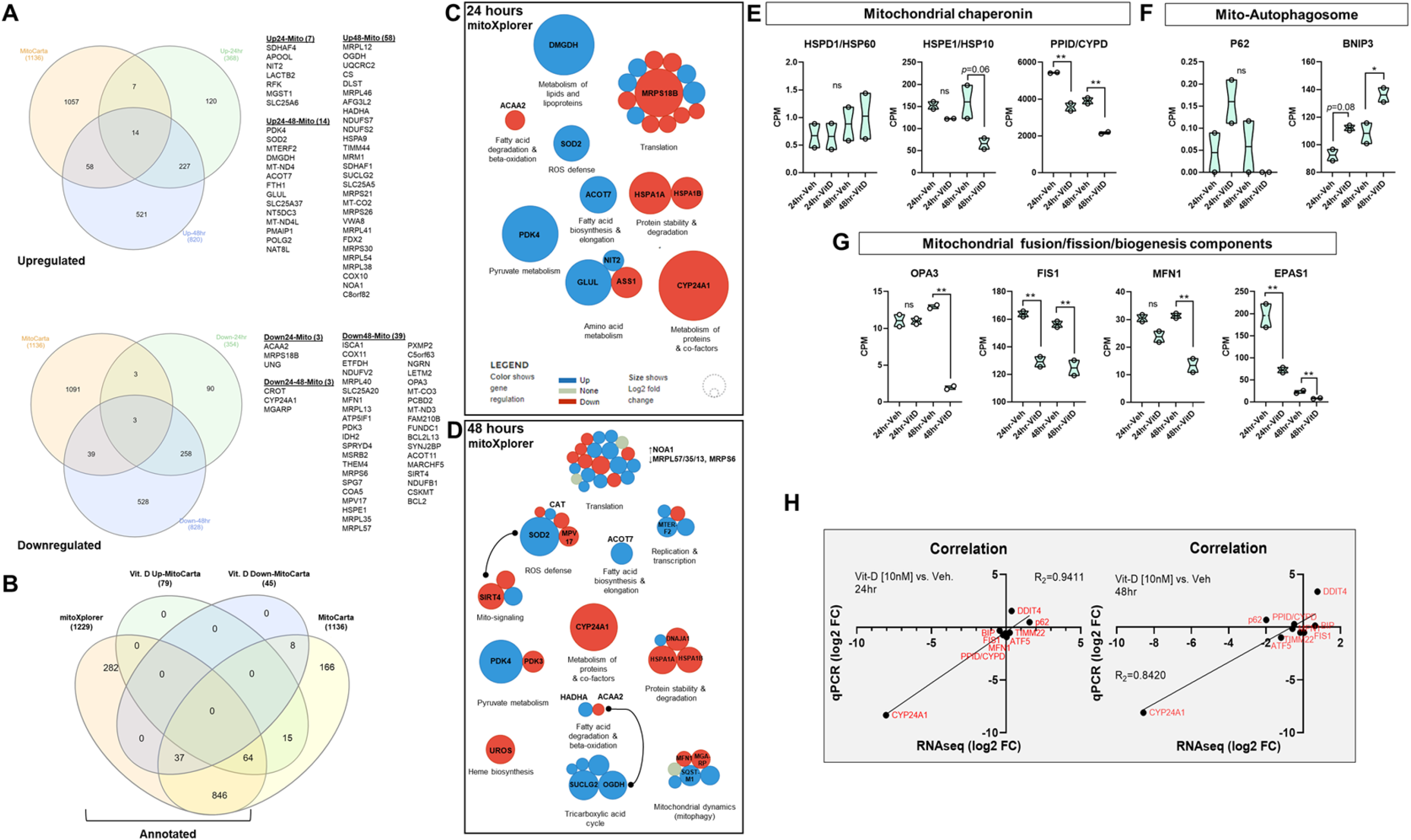
**A multi-omics approach to study mitochondrial anti-cancer responses to vitamin D** A) Identification of mitochondria-related genes from vitamin D treated MG-63 cells using MitoCarta. Differentially expressed genes (DEGs) from both the 24- and 48-hour datasets were cross-referenced to the MitoCarta database. Venn analysis was performed at http://www.interactivenn.net. B) Identification of annotated vitamin D-mediated mitochondrial genes. Mitochondrial DEGs derived from MitoCarta were cross-referenced with the annotated database mitoXplorer. Venn analysis was performed at http://www.interactivenn.net. C-D) Mitochondrial interactome of vitamin D treated MG-63 cells. Functional relationships were characterized by mitochondrial DEGs using the mitoXplorer software. http://mitoxplorer.ibdm.univ-mrs.fr/ E-G) RNAseq analysis of mitochondrial stress, biogenesis, and clearance regulators. A two-way ANOVA was performed with Bonferroni’s multiple comparisons test using the counts per million (CPM) values (n=2 samples/condition), where the *p*-value summaries were depicted as ^∗∗^*p* ≤ 0.01, and ^∗^*p* ≤ 0.05. ns (not significant) H) Real-time PCR validation of select RNAseq data. Plots showing correlation (R_2_= 0.94-0.84) between sample sets (n=3 samples/condition).

In the 48-hour analysis, the overwhelming effect of vitamin D on mitochondrial protein translation at 24 hours was aborted suggesting adaptive responses (**Figure 4D**). Additional selective pressures toward translation occurred via upregulation of MTERF2, a transcription termination factor that modulates cell growth and the cell cycle^32^. Longer treatments of vitamin D did enhance the ROS defense response (↑CAT); however, this was countered by decreased MPV17, which is involved in ROS neutralization and mitochondrial protection^33^. Antioxidant responses closely regulate mitochondrial epigenetic signaling factors such as SIRT4^34^, an enzyme with deacetylase and ADP-ribosylation activities, which was downregulated after vitamin D treatment, suggesting a mode for further fine-tuning of epigenomic regulation. Other mitochondrial metabolic and dynamic effects of vitamin D include the suppression of the heme biosynthesis pathway by way of UROS, which is part of the catalytic steps of porphyrin biosynthesis and associated with cancer when heme production is left unchecked^35^. Furthermore, mitofusion 1 (MFN1) was downregulated after vitamin D treatment that mediates mitochondrial fusion suggesting reduced mitochondrial networks, ATP production, and OXPHOS. SQSTM1, a protein involved in mitophagy, was upregulated after vitamin D treatment, suggesting a selective and adaptive process to remove dysfunctional mitochondria from cancer cells. The TCA cycle, which provides electrons via the reducing agent NADH for OXPHOS, was enhanced after 48 hours of vitamin D treatment despite the suppression of OXPHOS raising the possibility of non-redox roles^36^. For example, vitamin D may involve substrate-level phosphorylation as a metabolic reaction to generate energy instead of OXPHOS. With this in mind, SUCLG2, a GTP-specific beta subunit of succinyl-CoA synthase that forms succinyl-CoA, succinate, and ATP through the coupling of this reaction independent of OXPHOS, was elevated after vitamin D treatment. Also, OGDH, a dehydrogenase that catalyzes the conversion of 2-oxoglutarate to succinyl-CoA and carbon dioxide was increased after vitamin D treatment, which may further drive energy production via TCA non-redox intermediates.

Lastly, several known mitochondrial genes were not co-curated in the MitoCarta and mitoXplorer repositories including DDIT4/REDD1^37^ (see later) that were validated by qPCR derived from our RNAseq data sets (**Figure 4E-H**). We also observed a consistent downregulation of known mitochondrial chaperonin PPID (cyclophilin D). Although there was an upward trend for the mitophagy marker, P62 (**Figure 4F**), qPCR re-analysis showed a statistically significant increase in transcript levels (**Figure 4H**), suggesting a possible role in conjunction with SQSTM1 toward mitophagy. In addition, mitochondrial BCL2/adenovirus E1B 19 kDa protein-interacting protein 3 (BNIP3) transcripts were increased after vitamin D treatment and may interact with LC3 to remove damaged ER and mitochondria to recycle cellular content to promote the health of cells (**Figure 4F**). Again, in terms of mitochondrial dynamics, vitamin D treatment resulted in the down regulation of mitochondrial fission transcript, FIS1, and of OPA3, a dynamin-related GTPase that regulates the equilibrium between mitochondrial fusion and mitochondrial fission (**Figure 4G**). Endothelial PAS domain protein 1 (EPAS1) mRNA, which encodes a protein involved in mitochondrial biogenesis, was decreased after vitamin D treatment (**Figure 4G**). Overall, the multi-omics approach revealed novel factors and pathways as part of vitamin D’s mitochondrial-mediated anti-cancer response.

### Vitamin D-mediated epigenetic regulation of mitochondrial-related genes in MG-63 osteosarcoma cells

Next, to identify functional chromosomal regions that may govern anti-cancer responses and may be co-regulated by vitamin D and oxidative stress, we used <u>A</u>ssay for <u>T</u>ransposase- <u>A</u>ccessible <u>C</u>hromatin using <u>seq</u>uencing (ATACseq) and assessment of transcription factor (TF) binding motifs. This method appraises genome-wide chromatin accessibility using hyperactive Tn5 transposase that inserts sequencing adapters into open chromatin regions (**Figure 5A**). The data shows that most peaks were located within intronic, intergenic, and promoter regions across samples with 96-97% of reads with ≥Q30 scores, satisfying the quality control requirements (**Worksheet S9**). Globally, there were 97,739 overlapping peaks, 14,210 vitamin D-unique peaks, and 7,535 vehicle-unique peaks after vitamin D treatment for 24 hours (**Figure 5B,C**). The ATACseq results confirmed the RNAseq analysis showing an increased number of transcriptional start sites (TSS) that contain vitamin D-unique peaks at the expense of decreased nucleosome assembly (**Figure 5D, S2**). Because many *cis*-regulatory elements are close to the TSS of their targets, the data suggests that vitamin D promotes global chromatin accessibility, enrichment, and transcriptional regulation from the TSS. To get further insight, we compared key down and upregulated mitochondria-related transcripts identified with RNAseq to the list of significant peaks. For downregulated genes (**Figure 5E**), vitamin D treatment resulted in decreased chromatin accessibility at the distal promoter region of *ATF5*, suggesting possible regulation by negative vitamin D response elements^38^. *PPID* gene expression is directly regulated by ATF5, and we observed a similar decrease in chromatin accessibility at the TSS and proximal protomer region. Interestingly, *CYP24A1* was one of the most downregulated genes identified after vitamin D treatment, yet exhibited enhanced chromatin accessibility at both the TSS, proximal promoter, and as well as within intron 3-4, suggesting the possible “looping” of chromosomal structures that may suppress and discriminate CYP24A1 transcription in *trans* after vitamin D treatment^39^. For upregulated genes (**Figure 5F**), we observed enhanced chromatin accessibility at both the proximal and promoter regions of *DDIT4*. On the contrary, there appeared to be nominal epigenetic regulation of *SOD2* by vitamin D. This finding suggests either post-translational and/or post-transcriptional modes of SOD2 mRNA regulation after vitamin D treatment. Interestingly, one of the most significantly affected chromosomal regions identified by ATACseq was in intron 9-10 of the *SUCLG2* gene, suggesting a regulatory epigenetic mechanism induced by vitamin D to control succinyl-CoA synthase expression.

**Figure 5.**
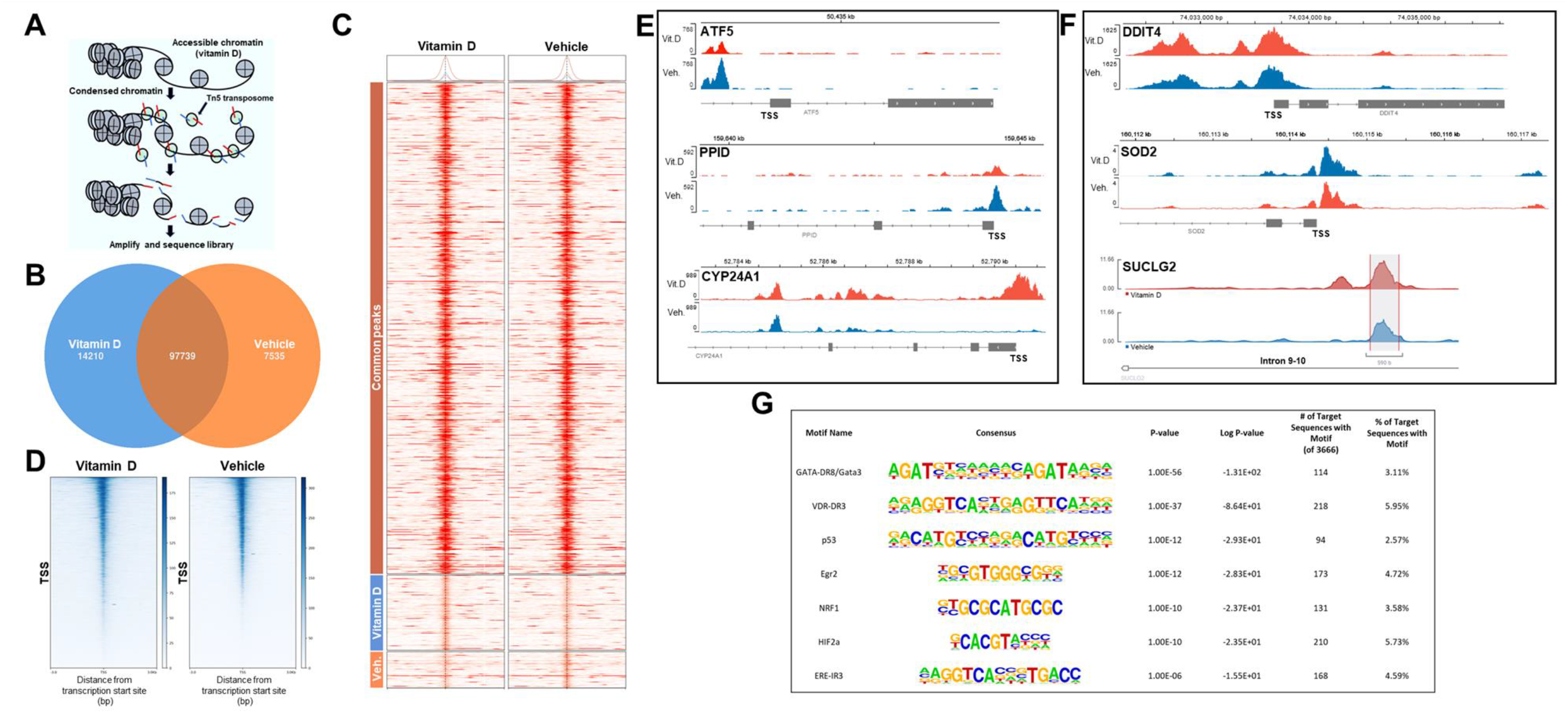
**VDR-mediated epigenetic regulation of MG-63 osteosarcoma cells** A) Changes in chromatin accessibility assayed by ATACseq. ATACseq identifies regions of open chromatin using Tn5 transposases and tagging sites with sequencing adaptors. B) Global peak overlaps and peaks unique to vitamin D [10nM] and vehicle treatments for 24 hours. C) Heatmap depicting the pattern of peaks outlined in the Venn diagram. There is one column per group used in the comparison. The color bars on the left correspond to Venn Diagram grouping of peaks. The heatmap displays the read coverage density (more red means more reads at that location), with each row corresponding to the average peak profile for a single peak that is averaged within each group, across all groups in the comparison. D) Heatmap for transcriptional start sites (TSS) after vitamin D [10nM] and vehicle treatments. Chromatin accessibility at TSS significantly increased in response to vitamin D. The lower half of the vitamin D plot harbors the vitamin D-unique peaks from B and C. The plots specify a window of +/- 3,000bp around the TSS of genes. Read density scores presented as the right index. E) Genome browser track of ATACseq results for select mitochondrial downregulated genes. Differentially accessible regions identified for select genes. Transcriptional start site (TSS) F) Genome browser track of ATACseq results for select mitochondrial upregulated genes. Of note, a vitamin D-unique peak was identified in intron 9-10 of *SUCLG2*. Transcriptional start site (TSS) G) List of transcription factor (TF) motifs enriched in accessible chromatin upon vitamin D treatment.

We next used Hypergeometric Optimization of Motif Enrichment (HOMER) for transcription factor (TF) motif discovery within vitamin D-sensitive open chromatin (**Worksheet S10**)^40^. From this analysis, GATA3 was the most highly associated TF identified (**Figure 5G**), whereby GATA3 is essential for normal tissue development^41^ and is commonly mutated in breast cancers^42^. Not surprisingly, the VDR motif was the second most highly correlated nuclear TF identified from our analysis. Interestingly, we also identified the nuclear respiratory factor 1 (NRF1), a redox-sensitive member of the Cap-N-Collar family of TFs that binds to antioxidant response elements (AREs)^43^, as a potential regulator of vitamin D-mediated epigenetic responses suggesting that AREs may cooperate with vitamin D response elements (VDREs) via NRF1-VDR binding. Vitamin D treatment also may promote estrogen receptor binding, suggesting a synergistic effect to help promote the normal bone-forming osteoblast phenotype in osteosarcoma cells^44^. Overall, vitamin D promotes chromatin accessibility in MG-63 cancer cells to enhance the regulatory effects of specific TFs that may play important roles in oxidative stress defense and normal tissue and cellular developmental processes.

### Vitamin D-mediated decrease in **ΔΨ_M_** inhibits mitochondrial ROS production

The transcriptomic and epigenomic data thus far suggests that vitamin D regulates mitochondrial functions in MG-63 cells to promote its anti-cancer effects. Therefore, we investigated the mitochondrial membrane potential (ΔΨ_M_) using the ratiometric JC-1 dye, where the accumulation of cationic J-aggregates (red) in mitochondrial membranes acts as a proxy for polarized mitochondria. On the other hand, cells that have diminished ΔΨ_M_ will contain JC-1 in its monomeric form (green) in either the mitochondria or cytoplasm during transition states. In the vitamin D studies, we pre-treated MG-63 cells for 24 hours and then measured the JC-1 intensity ratios (**Figure 6A**). Interestingly, while most of the vehicle-treated MG-63 cells were positive for J-aggregates, only ∼25% of vitamin D-treated cells contained J-aggregates in their mitochondria, suggesting a complete collapse of the ΔΨ_M_ within most cells (**Figure 6B**). Among those vitamin D-treated cells that exhibited J-aggregates, their JC-1 intensity ratio was significantly reduced compared to vehicle treatment (**Figure 6A,C**). Using the Imaris software (Bitplane) spot intensity tool, the vehicle-treated cells exhibited an overlap in J-aggregate-to-monomer signals across a series of mitochondria (i.e., spot-to-spot) within cells (**Figure 6D**). On the contrary, vitamin D-treated cells exhibited an increased level of non-overlapping monomer-to-J-aggregate signals, suggesting the depolarization of the mitochondria membrane and extramitochondrial presence of the monomers.

**Figure 6.**
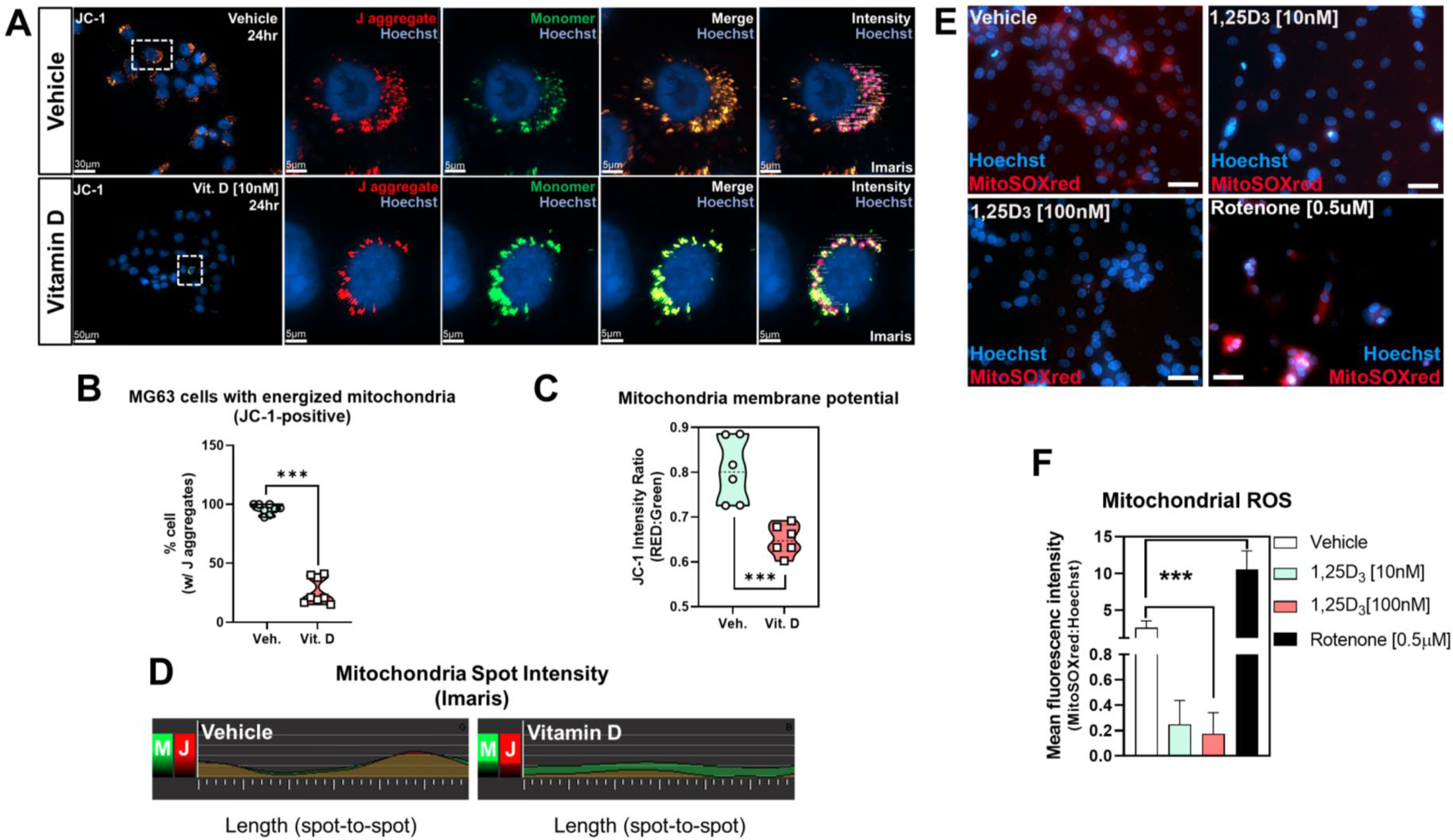
**Vitamin D depolarizes the mitochondrial membrane and inhibits ROS production** A). Mitochondria membrane potential (ΔΨ_M_) measured using the JC-1 cationic dye in MG-63 cells. B) Quantification of MG-63 cells with J aggregates after vitamin D [10nM] and vehicle treatments for 24 hours. Data are presented as mean±SEM error bars (n=8 replicates/condition); ^∗∗∗^*p* ≤ 0.001 (two-way ANOVA with Sidak’s multiple comparisons test compared to vehicle). C) Quantification of JC-1 intensity ratio in MG-63 cells treated with vitamin D [10nM] and vehicle for 24 hours. Data are presented as mean±SEM error bars (n=6 replicates/condition); ^∗∗∗^*p* ≤ 0.001 (two-way ANOVA with Sidak’s multiple comparisons test compared to vehicle). D) J aggregate and monomer dynamics in MG-63 cells. The spot intensity tool (Imaris) was used to determine the intensity (y-axis) and position (x-axis) between spot-to-spot (i.e., a mitochondria-to-mitochondria series) in treated cells. M (monomer), J (J aggregate). E) Mitochondrial superoxide detection in MG-63 cells after vitamin D [10nM] and vehicle treatments for 24 hours. Bar=20µm. F) Quantification of mitochondrial superoxide in MG-63 cells treated with vitamin D [10nM] and vehicle for 24 hours. Data are presented as mean±SEM error bars (n=6 replicates/condition); ^∗∗∗^*p* ≤ 0.001 (one-way ANOVA with Tukey’s multiple comparisons test compared to vehicle).

A key factor in determining the fate of cells with depolarized mitochondria is the level of ROS. To determine the impact of vitamin D on ROS production within MG-63 cells, we measured mitochondria-specific ROS using the MitoSOX™ Red, a mitochondrial O_2_ indicator for live cells (**Figure 6E**). Vitamin D treatment for 24 hours significantly reduced the production of O_2_ within MG-63 cells compared to vehicle-treated samples (**Figure 6F**). Conversely, treatment with the inhibitor of complex I of the respiratory chain, rotenone, significantly increased mt ROS levels. Thus, MG-63 osteosarcomas are accompanied by mechanisms that prevent mitochondrial depolarization, resulting in chronic intracellular mt ROS. Overall, the data suggest that vitamin D treatment is associated with the opening of the mitochondrial permeability pores, loss of the electrochemical proton gradient and reduced ROS as part of its anti-cancer effects.

### Vitamin D modulates mitochondrial structure and dynamics in MG-63 cancer cells

We next appraised mitochondria structure and morphology using immunofluorescence (IF) and electron microscopy (EM). Using antibodies against the outer mitochondrial membrane voltage-dependent anion-selective channel 1 (VDAC-1), we observed the classic elongated tubular shape of mitochondrial in vehicle-treated MG-63 cells (**Figure 7A**). However, vitamin D treatment promoted changes from a tubular to globular “ring” morphology with weak fluorescence in the center of mitochondria (**Figure 7B**). This characteristic suggests mitochondrial shortening, swelling, and structural changes trigged by decreased ΔΨm and increased permeability of the inner mitochondrial membrane. 3D-rendered images of vitamin D-treated cells revealed rough, fragmented surfaces (i.e., structural changes) of individual mitochondria treated for 24 hours (**Figure 7B, lower panel**). The reversible changes of mitochondria from the tubular to condensed or fragmented conformations is the classic response of loss of ATP synthesis by OXPHOS^45^.

**Figure 7.**
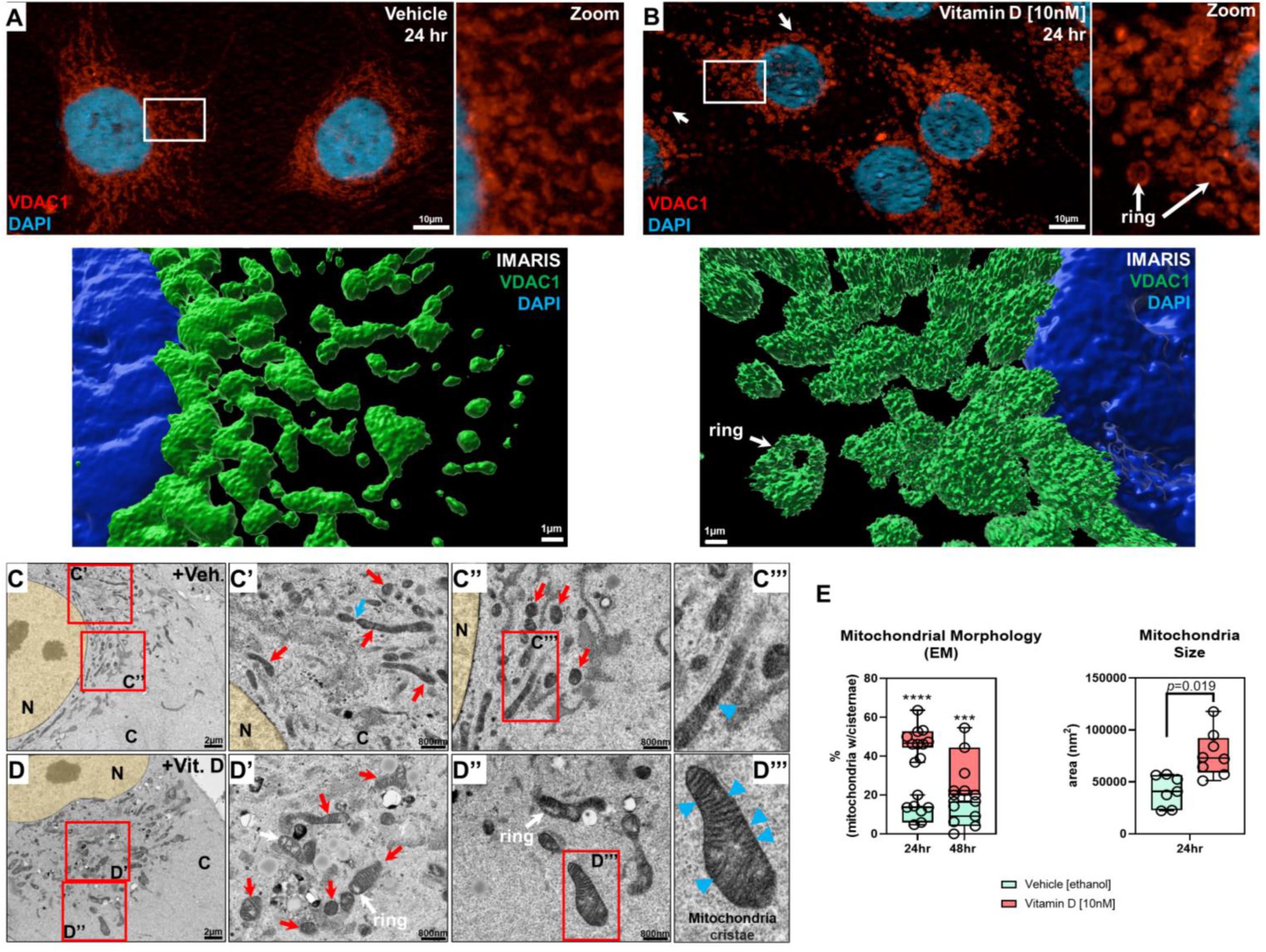
**Vitamin D modulates mitochondria structure in MG-63 osteosarcoma cells** A) Immunofluorescence labeling of VDAC1 within vehicle-treated MG-63 cells. Right panel is the magnification of the inset. The lower panel is Imaris 3D rendered image of the inset. B) Immunofluorescence labeling of VDAC1 within vitamin D [10nM] treated MG-63 cells. The right panel is the magnification of the inset. Arrows depict mitochondrial ring-like structures. The lower panel is Imaris 3D rendered image of the inset. C, C’, C”, C””) Representative transmission electron microscopy (TEM) images of vehicle- treated MG-63 cells for 24 hours. Red insets are marked with panel identifiers. C’, blue arrow depicts tethered mitochondria, and red arrows depict tubular mitochondria. C”, red arrows depict electron-dense cross sections of tubular mitochondria. C””, blue arrowhead depicts loosely structured cristae. C (cytoplasm), N (nucleus). D, D’, D”, D””) Representative TEM images of vitamin D [10nM] treated MG-63 cells for 24 hours. Red insets are marked with panel identifiers. D’, red arrows depict mitochondria in various stages, e.g., tubular, herniated, swollen, with visible cristae. White arrows depict rings in mitochondria. D””, blue arrowheads depict defined cristae structures in mitochondria. E) Quantification of TEM. For analysis, between 7-10 cells were investigated per condition, in which we averaged parameters between 20-40 mitochondria per cell. Data are presented as mean±SEM error bars (n=7-10 cells/condition);); ^∗∗∗∗^*p* ≤ 0.0001, ^∗∗∗^*p* ≤ 0.001 (two-way ANOVA with Bonferroni’s multiple comparisons test compared to vehicle).

In EM studies, vehicle-treated cells exhibited tubular mitochondria, however individual cristae were hardly discernable, suggestive of deranged mitochondrial respiration^46^ (**Figure 7C, red arrows**). Furthermore, some mitochondria were in various stages of membrane fusion/fission as marked by “tethered” structures indicative of dynamic remodeling (**Figure 7C, blue arrow**). Vitamin D treatment increased the size of mitochondria and generated mitochondria with discernable cristae (**Figure 7D,E**), which may reflect partial prevention of cristolysis. The mitochondria also contained electron-lucent cavities, not vacuoles^47^, consistent with the ring-shaped structures observed in the IF studies (**Figure 7B**, white arrow) and may represent interorganellar connections commonly observed in normal cells^48^.

### Vitamin D regulation of mitochondrial biogenesis mediates DDIT4/REDD1 availability and mTOR function in the cytoplasm

Lastly, given the results of our functional annotation analysis and recent findings that certain cells express DDIT4/REDD1 in the mitochondria^49^, we focused the remainder of our attention on the role that vitamin D and DDIT4 play in cancer prevention. DDIT4 is a known tumor suppressor gene predominantly expressed in the cytoplasm under certain stress conditions to function as a potent mTOR inhibitor^50^. However, recent findings show that DDIT4 is highly expressed in malignant cancers leading to poor cancer-related prognosis in a paradoxical manner^18, 37^, suggesting that for certain genes the expression profiles cannot be functionally generalized (**Figure S3**). To help rationalize this paradoxical observation, we investigated DDIT4 cellular flux in MG-63 cells before and after vitamin D treatment. First, vitamin D at 10nM increased DDIT4 mRNA levels in a time-and VDR-dependent manner (**Figure 8A**). Next, we performed Apotome (Zeiss) structured-illumination imaging of DDIT4 and VDAC1 within vehicle-treated MG-63 cells and found that DDIT4 was exclusively associated with VDAC1-positive mitochondria (**Figure 8B,** yellow arrow). Imaris 3D-rendering confirmed the proximity of both VDAC1 and DDIT4 within individual mitochondria (**Figure 8C**), whereby VDAC1-free DDIT4 expression in the cytoplasm was uncommon (white arrow) (**Figure 8C**). A more dynamic colocalization pattern was observed after vitamin D treatment (**Figure 8D,E**). DDIT4 was predominantly expressed in the cytoplasm after vitamin D treatment (**Figure 8E**). Interestingly, among the VDAC1-DDIT4 co-localized mitochondria in vitamin D-treated cells, the average spot distance (i.e., the shortest distance between VDAC1 and DDIT4) was significantly increased compared to controls suggesting the translocation or leaking of DDIT4 into the cytoplasm (**Figure 8F, bottom panel**). Furthermore, after rotenone treatment, MG-63 cells contained globular VDAC1-positive mitochondria as previously noted, as well as a disassociation of DDIT4 from the mitochondria, suggesting a potential role of mitochondrial depolarization (**Figure 8H**). Nonetheless, the level of co-localized VDAC1-DDIT4 protein in vitamin D-treated cells was significantly decreased compared to vehicle treatment, reflecting the excess of DDIT4 in the cytoplasm (**Figure 8G**). In contrast, vehicle-treated MG-63 cells contained a statistically significant higher percentage of VDAC1-DDIT4 colocalized mitochondria with lower levels of cytoplasmic DDIT4 (**Figure 8G**). Given the apparent decrease in the number of mitochondria after vitamin D treatment, the increase in cytoplasmic DDIT4 protein may occur, in part, due to reduced mitochondrial biogenesis (i.e., mass/content/self-replication). To address this possibility, we used a duplexing in-cell ELISA assay that quantifies both mitochondrial (mt) DNA- and nuclear (n) DNA-encoded proteins, COX-1 and SDHA, respectively, in MG-63 cells. We observed vitamin D dependent inhibition of mtDNA-encoded COX-1 protein relative to nDNA-encoded SDHA protein by ∼20% after 24 hours (**Figure 8I**). These data suggest that enhanced mitochondrial localization of DDIT4 may help confer the cancer state and that the enhanced cytoplasmic localization and expression of DDIT4 may be a mechanism by which vitamin D suppresses osteosarcomas.

**Figure 8.**
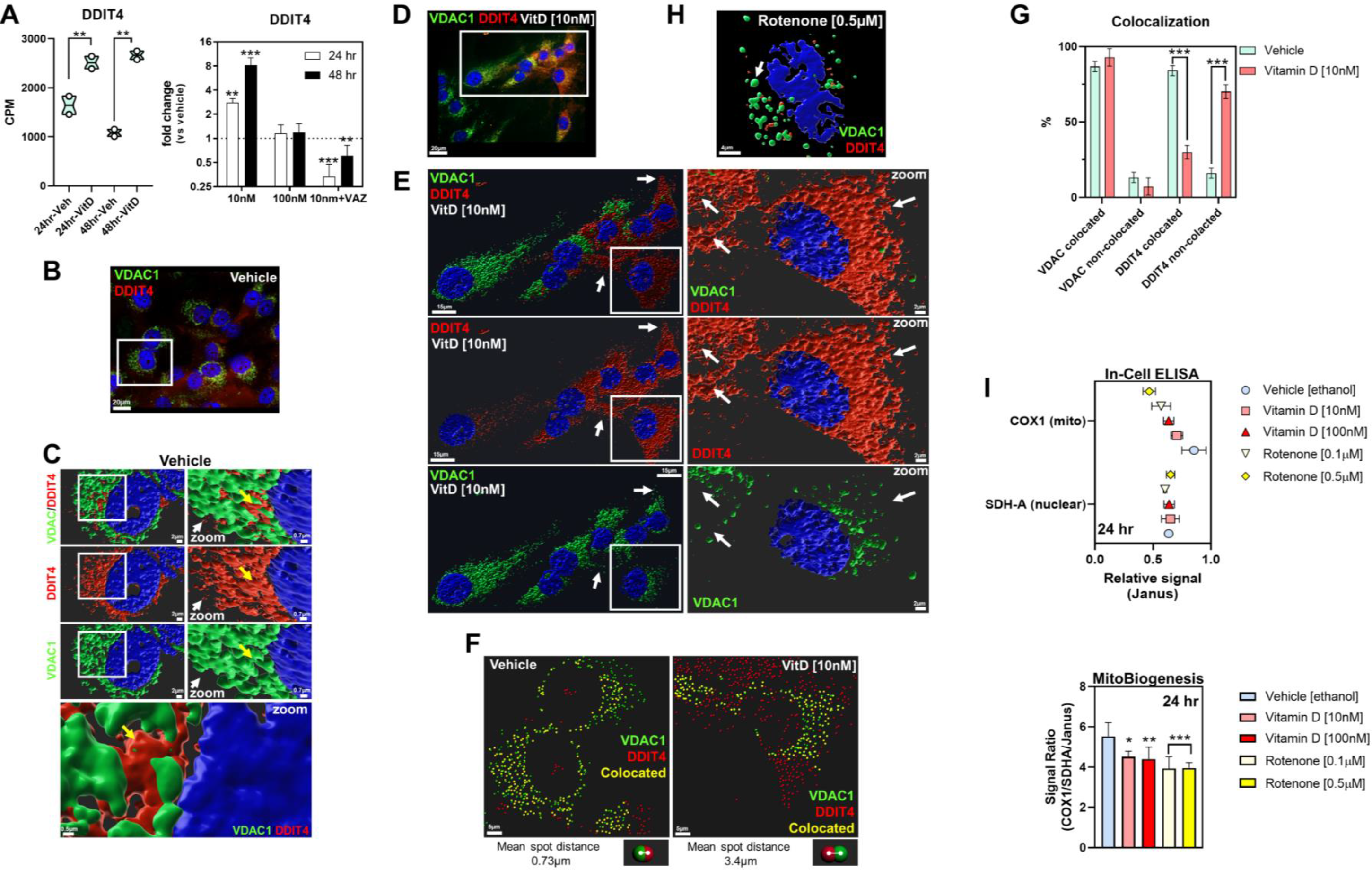
**Vitamin D regulation of mitochondrial biogenesis and DDIT4/REDD1 cytoplasmic availability** A) DDIT4 transcript levels after vitamin D treatment of MG-63 cells. The left panel depicts the RNAseq data whereby a two-way ANOVA was performed with Bonferroni’s multiple comparisons test using the counts per million (CPM) values (n=2 samples/condition). The *p*- value summaries are depicted as ^∗∗^*p* ≤ 0.01. The right panel depicts the real-time PCR validation at two different durations and vitamin D concentrations. VAZ was used as a VDR- specific inhibitor. The data are presented as mean±SEM error bars (n=3 samples/condition); ^∗∗∗^*p* ≤ 0.001, and ^∗∗^*p* ≤ 0.01 (one-way ANOVA with Tukey’s multiple comparisons test compared to respective vehicle). B) Immunofluorescence labeling of vehicle-treated MG-63 cells for VDAC1 and DDIT4 after 24 hours. C) 3D Imaris rendering of inset in B. The right panel depicts the magnification of the left panel inset. Yellow arrows depict VDAC1-DDIT4 co-localized mitochondria, while the white arrow depicts sparse cytoplasmic DDIT4. The lower panel depicts high magnification of VDAC1-DDIT4 colocalization. D) Immunofluorescence labeling of vitamin D treated MG-63 cells for VDAC1 and DDIT4 after 24 hours. E) 3D Imaris rendering of inset in D. The right panel depicts the magnification of the left panel inset. White arrows focus on positions of DDIT4 expression relative to VDAC1 placement. F) Representative image of VDAC1-DDIT4 colocalization and separation after vitamin D treatment for 24 hours using Imaris. Co-localization and separation analysis was performed using the Imaris “spot” tool to designate the distance threshold and the mean distance between the “VDAC” and “DDIT4” co-localized spots. Yellow spots depict colocated elements. Bottom panel depicts the shortest mean spot distances for each treatment conditions across all colocated spots. G) Quantification of VDAC1 cocoalization after vitamin D treatment for 24 hours using Imaris. A two-way ANOVA test with Sidak’s multiple comparisons test was performed between vehicle and treatment data sets using Prism (GraphPad) where the *p*-value summaries were depicted as ^∗∗∗^*p* ≤ 0.001. Statistical significance was accepted at ^∗^*p* ≤ 0.05. I) Mitochondrial biogenesis and translation assay after vitamin D treatment for 24 hours in MG- 63 cells. The upper panel depicts the relative signal of COX1 and SDH-A normalized to Janus. The bottom panel depicts the level of mitochondrial biogenesis and translation based on the signal ratio of measured factors. Data are presented as mean±SEM error bars (n=5 replicates/condition); ^∗∗∗^*p* ≤ 0.001, ^∗∗^*p* ≤ 0.01, ^∗^*p* ≤ 0.05 (two-way ANOVA with Tukey’s multiple comparisons test compared to vehicle).

## Discussion

### Relationship between vitamin D and the metabolic oxidation/reduction reactions of cancerous and non-cancerous cells

Findings so far in non-cancerous cells suggest that proper vitamin D levels maintain and minimize systemic cellular oxidative stress following the day-to-day exposure to damaging agents such as UV sunlight^51^. Furthermore, loss of VDR functional studies in human skin keratinocytes show increased mitochondrial membrane potential due to increased transcription of the respiratory chain subunits II and IV of cytochrome c oxidase^52^. In addition, the potential for vitamin D to reduce oxidative damage to DNA has been linked to a clinical trial where vitamin D supplementation reduced 8-hydroxy-2′-deoxyguanosine, a marker of oxidative damage, in colorectal epithelial crypt cells^53^. In other studies, vitamin D was shown to modulate the expression of select anti-oxidative genes via nuclear factor erythroid 2-related factor 2 (NRF2), which is a key transcription factor that can bind to AREs to protect cells against oxidative stress associated with diabetic neuropathy^54^. These findings suggest that vitamin D can regulate the respiratory chain and to modulate ancillary metabolic pathways depending on the cellular context and requirements within stressed non-cancerous cells.

Our findings in cancer cells show that vitamin D can influence mitochondrial metabolism, structure, and function to dictate its anti-cancer effects which may also intimately involve extra- mitochondrial organelles such as the ER (**Figure 3**,**9**). Membrane potential is directly related to the activity of mitochondria, with more activity correlated with higher stress levels. Our findings show that there is lower mitochondria activity through the depolarization of the mitochondrial membrane after vitamin D treatment, hence less stress and ROS production. Vitamin D decreased the mitochondrial membrane potential to a level sufficient for cells to survive but insufficient to generate mt ROS, unlike other mitochondrial depolarizers such as hydrogen peroxide^55, 56^. Mitochondrial depolarization aims to reverse the inner membrane potential to generate minimal ATP with lower mt ROS, which is associated with increased longevity as observed in long-lived naked mole rats and bats^57^. By contrast, in the absence of vitamin D, MG-63 osteosarcomas are accompanied by mechanisms that prevent mitochondrial depolarization, resulting in chronic intracellular protein and DNA damage by mt ROS^58^. In addition, vitamin D treatment resulted in mitochondria with “ring-like” structures, which may represent intracellular organellar connections such as mitochondria-ER associated membranes where the ER has been pulled into the lumen of the mitochondria to facilitate the direct transfer of phospholipids between the ER and mitochondria to help organize and generate cristae^48^. It is also possible that vitamin D may promote the sharing of other interorganellar resources such as mitochondria-derived antioxidants or SOD2, to support the stressed ER, that has the potential to produce additional ROS itself. In contrast, untreated MG-63 cells exhibit a lack of distinct mitochondrial cristae suggesting defects in the conductivity of structural proteins and stages of replication with constant flux between fusion and fission where the double-membrane structures are not as well developed. The potential reduction in fusion and fission after vitamin D treatment may also result in reduced complementation of mitochondrial gene products that occurs to repair damaged mitochondria in untreated cancer cells. That is to say, the reduced fission/fusion may reflect the limited need for organellar quality control, supported by the downregulation of ATF5. Overall, in MG-63 cancer cells, vitamin D functions to decrease mt ROS levels, which may also prevent ROS leakage to other cellular compartments to maintain molecular and organellar integrity.

**Figure 9.**
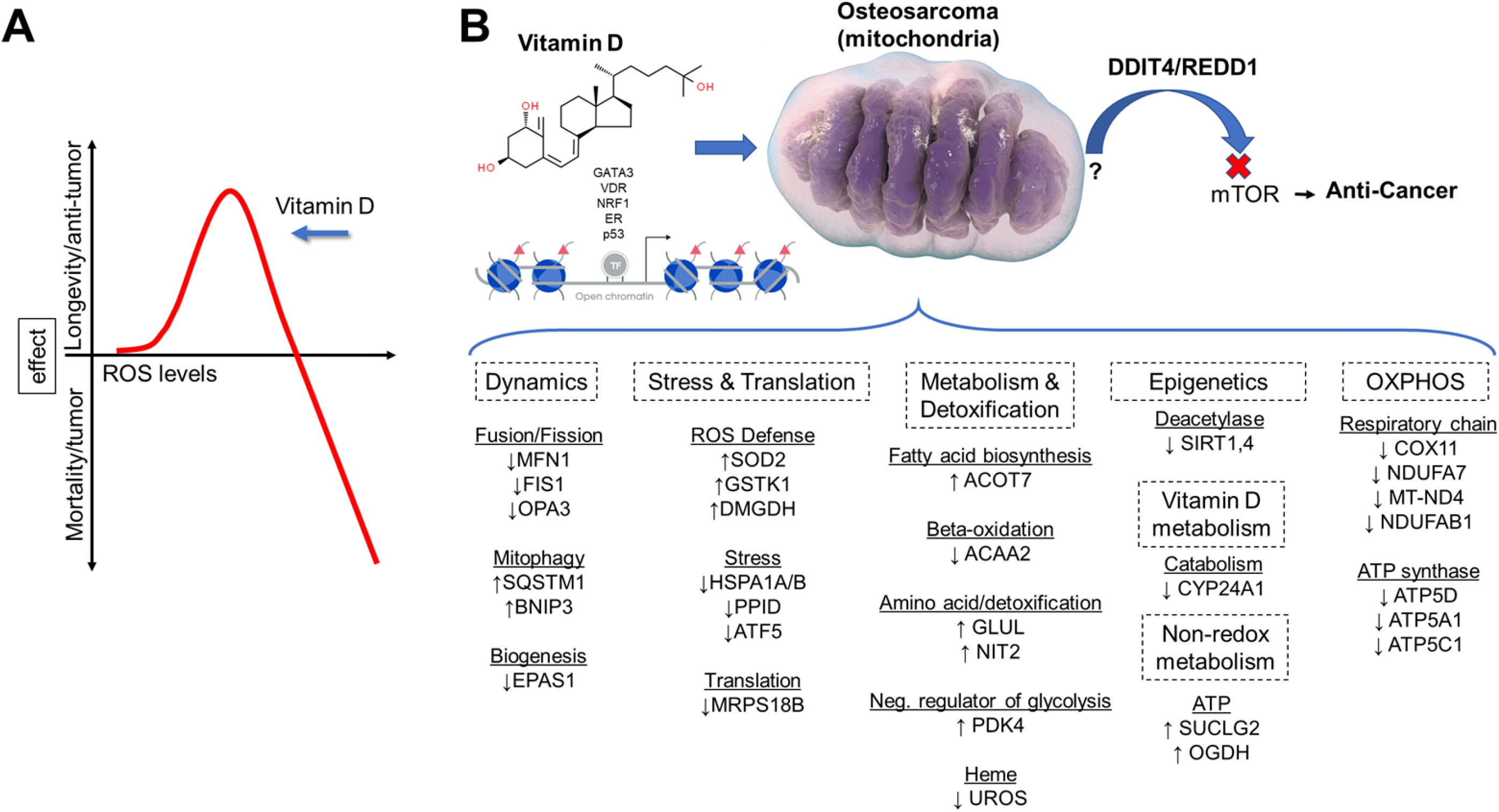
**Vitamin D regulation of mitochondrial hormesis and functions within MG-63 osteosarcoma cells** A) Vitamin D suppresses cancer progression by modulating the ROS threshold. Vitamin D- mediated mitohormesis and ROS production reprograms MG-63 osteosarcoma cells to enforce stress tolerance and growth inhibition to improve health outcomes. B) Vitamin D modulates chromatin assembly and mitochondrial metabolism, oxidative stress and mTOR inhibition via DDIT4/REDD1 localization to dictate its overall anti-cancer response. Red triangles represent acetylation.

### Vitamin D and ROS regulation

How vitamin D reduces mitochondrial ROS levels is unclear, and likely involves multiple factors including cellular detoxification as well as reduced synthesis in the mitochondria via downregulation of the respiratory chain complex subunits. Although we showed that vitamin D can enhance SOD2 levels, there are other ROS scavenging and degradation systems such as glutathione peroxidase, glutathione reductase, thioredoxin, and catalase, some of which were also regulated by vitamin D treatment in our studies. Our results also show that vitamin D can decrease the beta oxidation of fatty acids as an additional means to reduce ROS levels. Vitamin D may also regulate ROS production and turnover in other organelles besides mitochondria. ROS are also produced in the ER through the oxidation of proteins. Although our ER stress results show the activation of the early stage of the UPR after vitamin D treatment, the more severe consequences of unfolded proteins (e.g., ER-mediated apoptosis) were not evident, suggesting that vitamin D may also prevent this from occurring. In addition, vitamin D induced heme oxygenase-1 (HMOX1) expression, which is an essential enzyme localized to the plasma membrane and Golgi apparatus for heme catabolism to form biliverdin, carbon monoxide and ferrous iron^59^. Free heme promotes ROS production by damaging DNA, lipids, and proteins. In conjunction, the generation of carbon monoxide may also regulate key anti-inflammatory cytokines (IL-10, IL-1RA, ↑<u>NFKBIA</u>) that may play an anti-cancer role in MG-63 cells after vitamin D treatment^6^.

### Vitamin D and ROS consequences

The direct consequences of lowered ROS levels after vitamin D treatment on intracellular targets and processes in cancer cells are unknown. It is known that at low levels, ROS can act as signaling molecules in various intracellular processes such as migration. At low levels, ROS differentially alters target protein conformation to modulate stability, folding, and activities of enzymes and ensuing phosphorylation cascades. The decrease in ROS production mediated by vitamin D is likely to also affect downstream cystine oxidation of proteins to regulate, for example, DNA repair and damage, to facility the anti-cancer response. Overall, our results suggest that after vitamin D treatment the mitochondria produce low levels of ROS as a new set point that are effectively scavenged by the cancer cells’ antioxidant defense system. It is at this new set point that makes mitochondrial ROS an intracellular signaling molecule that does not induce oxidative stress, instead provides a safe “basal” window for redox signaling to alter the cancer cell fate.

Cell studies have shown that ROS can have both inductive and suppressive effects on global transcription as well as epigenetic responses in a highly cell-type specific manner^60^. This may be related to the variable epigenetic and ROS-sensitive and/or insensitive transcription factors, or factors that control the availability of antioxidants thiols within a cell. The effects of ROS on epigenetic and genetic mechanisms can involve direct effects that entail modifications of DNA bases and histones, or indirect effects on DNA and histone-modifying enzymes to control cancer development^61^. ROS can also affect class III histone deacetylases (HDACs) called sirtuins (SIRTs). Unlike class I, II, and IV HDACs that use metals as cofactors, SIRTs require NAD^+^ as a functional cofactor thus making this enzyme sensitive to metabolic and redox changes to relay cellular stress in the form of histone modifications and changes in gene expression^62^. In our study, vitamin D treatment resulted in the downregulation of the mitochondrial SIRT1/4 deacetylases in MG-63 cells, which may signify a decoupling of lysine deacetylation with NAD+ hydrolysis and PDK4-acetly-CoA (histone acetylation) to promote gene expression. Tumor studies have shown that SIRT4 has both oncogenic and tumor-suppressive activities in cancer depending on the experimental conditions^63^. In the context of vitamin D signaling and concomitant ROS reduction, SIRT1/4 downregulation may help create an epigenomic landscape and balance to facilitate vitamin D-specific anti-cancer transcriptional responses and genomic stability.

### Vitamin D and stress tolerance and metabolic responses

Unchallenged protein misfolding can elicit cell death, while low levels of stress may be beneficial to cells by eliciting an adaptive UPR^23^. Furthermore, the beneficial effects of mild stress on aging and longevity have been studied in experimental animals, whereby mild dietary stress by way of dietary restriction without malnutrition delays age-related physiological changes and extends the lifespan. Importantly, animal studies have also demonstrated that mild dietary stress can prevent or lessen the severity of cancer^64^. Recent findings using the model organism, *Caenorhabditis elegans*, showed that vitamin D can promote longevity by enhancing proteostasis^65^, which may be akin to our findings of mitochondrial proteostasis and reduced biogenesis in MG-63 cells. These findings suggest that vitamin D may mimic a metabolic state induced by dietary restriction and/or mild UPR to improve the lifespan and anti-cancer effects.

Indeed, our previous studies showed that vitamin D treatment was comparable to serum starvation of cultured osteoblasts, where suppression of the mTOR pathway was identified as a common feature and known to also be involved in lifespan expansion in mice when inhibited with rapamycin^66^. Furthermore, our RNAseq and ATACseq motif analysis revealed associations with hypoxia, suggesting that vitamin D may promote tumor starvation by inhibiting vascular perfusion less the negative effects of elevated ROS. Also, vitamin D can promote mitochondrial depolarization, which is coupled to the availability of glucose or creatine, akin to dietary restriction to support sufficient mitochondrial ATP. These observations can also be metabolically linked to the increase in PDK4 we observed after vitamin D treatment. PDK4 is increased during hibernation/starvation and helps to decrease metabolism and conserve glucose by reducing its conversion to acetyl-CoA for ATP production^67^.

Our model suggests that vitamin D changes the metabolism of cancer cells from being responsive to stress to that of tolerant of stress that involves ER/mitohormetic processes with overall ROS reduction (**Figure 3**,**9**). There is recent precedence for this model in the natural immunometabolism setting involving microbial-macrophage interactions^68^. Timblin et al. showed that modulation of initial elevated antimicrobial ROS levels within macrophages involves ROS defense strategies as well as metabolic shifts toward non-oxidative energy metabolism resulting in a reduction of ROS levels for macrophages to survive and function. Our model similarly shows a parallel paradigm enforced by vitamin D on the dysregulated metabolism of MG-63 cancer cells. Co-opting this stress tolerance response identified in this study by vitamin D may be a future strategy to consider toward cancer therapy. Importantly, we identified key vitamin D-mediated metabolic enzymes that regulate fluxes of small compounds to provide the appropriate basal substrates for cell structure and energy production within dysfunctional osteosarcoma cells. For example, vitamin D upregulated DMGDH, whereby it acts as an antioxidant when its enzymatic byproduct, dimethylglycine, is used to support the one-carbon (1-C) metabolism toward cytosolic NADPH production^28^. Importantly, increased DMGDH levels are linked to hepatocellular carcinoma suppression^29^. Furthermore, vitamin D also positively regulates succinyl-CoA synthase which facilitates the coupling of succinyl-CoA synthesis and hydrolysis to substrate level phosphorylation of ADP to ATP^36^. The significance of this finding is that despite mitochondrial depolarization and OXPHOS inhibition after vitamin D treatment, the cell can generate sufficient ATP via non-redox metabolism independent of mitochondrial electron acceptors to support anti-cancer biological activities, including survival.

### Linking vitamin D regulation of DDIT4/REDD1 to mitochondria and cancer biology

In the physiological setting, DDIT4 is highly expressed in the cell cytoplasm under stress conditions such as hypoxia, cigarette smoke^69^, and UV-induced DNA damage to function as a potent mTOR inhibitor to suppress cell proliferation and growth, while promoting autophagic processes instead. DDIT4 is also highly expressed in malignant cancers^18, 37^, despite its known mTOR-inhibiting properties, suggesting that some cancers have evolved mechanisms to resist DDIT4, which may also antagonize anti-tumor therapies. For example, a meta-analysis of individual cancer datasets using Gene Expression Profiling Interactive Analysis (GEPIA) shows that DDIT4 mRNA expression is significantly increased in numerous tumor tissues such as cervical squamous cell carcinoma (CESC)^18^ (**Figure S3**), however, no data on osteosarcoma is currently available. We use GEPIA to further determine the overall cancer survival for CESC based on *DDIT4* gene expression levels. DDIT4 levels were normalized for relative comparison between a housekeeping gene, *ACTB*, and the *VDR* gene. Using the log-rank test (Mantel-Cox test) for hypothesis evaluation, the hazard ratio (HR) and the 95% confidence interval information associated with both gene normalization comparisons suggest a significant association with decreased survival of patients with elevated DDIT4 levels (*p*=0.0019&0.039 and HRs 2.1&1.6). The *VDR* relative comparison resulted in a higher *p*-value and lower HR, suggesting direct regulation of DDIT4 levels by vitamin D across individuals. This association of decreased survival for high DDIT4 cohorts was observed for many other cancer types besides CESC presented in GEPIA, suggesting elevated DDIT4 is associated with poor prognosis and a vitamin D component.

In line with the findings from GEPIA, our findings in MG-63 cancer cells show that the mitochondria and their biogenic state can dictate DDIT4 cellular localization pattern and function. In contrast to MG-63 cancer cells, our previous findings using normal primary osteoblasts showed a robust cytoplasmic expression pattern of DDIT4 under basal settings^17^, which suggests a DDIT4 dichotomy between normal and cancer states. Currently, it is unknown if DDIT4 mitochondrial sequestration and biogenesis are a generalized feature of most cancer cell types, and it is likewise unknown how vitamin D can regulate DDIT4 organellar sequestration and functional outcomes in those cancer cell types. Interestingly, we used an *in silico* mitochondria targeting sequence (MTS) predictor and identified a putative MTS only in the n-terminus of DDIT4 that contains a cysteine residue in the cleavage domain (**Figure S3**).This suggests that vitamin D, through its effects on ROS production, may regulate DDIT4 interactions via reactive cysteines with the mitochondria. Given the unknown function of DDIT4 in the mitochondria of MG-63 cells, future studies will focus on better understanding its role in the regulation of cell metabolism and mitochondrial biogenesis in the context of vitamin D treatment and oxidative signaling.

## Author contributions

T.S.L. conceived, designed, and performed the experiments, analyzed the data, and wrote the manuscript. Intellectual contributions were made by all coauthors. M.Q. performed the experiments, analyzed the data, wrote part and edited the manuscript. E.C., Z.W., H.Z., M.H., and S.R. analyzed the data and edited the manuscript. All authors gave final approval to the manuscript.

## Acknowledgment

Special acknowledgments to Karen Di Lauro, Chaitanya Doshi and Neda Vishlaghi (University of Miami) for providing technical support. We would like to thank Vania Almeida for assistance with transmission electron microscopy (U. Miami, Miami Project to Cure Paralysis) and Marissa Brooks with RNA and ATAC sequencing (U. Miami, Sylvester Comprehensive Cancer Center). Supported by Grant # IRG-17-183-16 from the American Cancer Society, and from the Sylvester Comprehensive Cancer Center at the Miller School of Medicine, University of Miami

## Data availability statement

All data generated during and/or analyzed during the current study are available from the corresponding author on reasonable request.

## Experimental procedures

### Reagents and cell culture

Crystalline 1,25(OH)_2_D_3_ (Millipore Sigma, 679101) and the vitamin D receptor antagonist ZK159222 (VAZ, Toronto Research Chemicals) were reconstituted in ethanol and kept at -80°C. Human MG-63 osteosarcoma cells (CRL-1427; American Type Culture Collection, Manassas, VA, USA) were cultured in complete media containing Eagle’s Minimum Essential Medium (ATCC, 30-2003), 10% heat-inactivated fetal bovine serum (Gibco), and 100 U/mL penicillin, 100 mg/mL streptomycin (Life Technologies). For assays, cells were treated with 0 (vehicle; equal-volume ethanol; 0.0001%), 10nM and 100nM 1,25(OH)_2_D_3_ incubated in tissue culture plates (CytoOne) at 37°C in a humidified atmosphere of 5% CO_2_, 95% air.

### Soft agar colony formation assay

MG-63 cells (1000 per well, 24-well plate) were seeded into 0.4% low melting point agarose (Lonza, 50101) on top of a 1% agarose layer. Cells were maintained in a 5% CO_2_ incubator at 37°C for approximately 14 days with vehicle or vitamin D. Colonies were fixed in methanol and stained with crystal violet. For quantification, crystal violet-positive colonies were counted using a dissecting scope (Zeiss Stereo 305) with the ImageJ software. All assays were set up in five-six replicates per condition. A one-way ANOVA test was performed with Tukey’s multiple comparisons test.

### RNA sequencing and functional/pathway/gene set enrichment analyses

Cell preparations from two independent experiments were collected and total RNA was purified using the PureLink RNA Mini kit (ThermoFisher Scientific, 12183018A) with DNase set (ThermoFisher Scientific, 12185010). RNA quality was tested by Agilent Bioanalyzer and confirmed to have RIN numbers > 8.5. Library preparation and RNA-sequencing were performed at the Oncogenomics Core Facility at the Sylvester Comprehensive Cancer Center (U. Miami). Samples were sequenced using 75bp paired ends with an Illumina NextSeq 500 generating reads of ∼30 million per sample that were trimmed and filtered using Cutadapt. Compressed Fastq.gz files were uploaded to a Galaxy account (https://usegalaxy.org/)^70^, and datasets were concatenated tail-to-head (Galaxy Version 0.1.0). A MultiQC analysis was performed on all FastQC raw data files to generate a summarized QC report. HISAT2 was performed for sample alignment to the human genome (Ensembl 87: GRCh38.p7 human transcriptome) to generate BAM files with the most reads (80–90%) aligning successfully. Samstools stat was performed for further quality control of BAM files, and HTseq-count was performed to generate non-normalized gene counts for each sample. Normalization of RNAseq gene counts (counts per million, CPM), and differential gene expression and visualization analyses were performed with iDEP (http://bioinformatics.sdstate.edu/idep93). Differentially expressed genes were determined using DESeq2 with the false discovery rate (FDR) set to 0.05 and logFC(1) compared to controls. For individual gene expression plots derived from RNAseq, a two-way ANOVA was performed with Bonferroni’s multiple comparisons test using the CPM values, where the *p*-value summaries were depicted as ^∗∗∗∗^*p* ≤ 0.0001, ^∗∗∗^*p* ≤ 0.001, ^∗∗^*p* ≤ 0.01, and ^∗^*p* ≤ 0.05. Gprofiler was utilized to perform gene name conversion (ENTREX TO ENTRZ), and basic functional annotation analyses (http://biit.cs.ut.ee/gprofiler/gost). To adjust for the FDR, we only considered terms with a Benjamini–Hochberg adjusted *p*-value of 0.05. iDEP was utilized for the distribution of transformed data and the generation of scatter plots of sample correlations, hierarchical and k-means heatmap generation, and pathway analyses. GSEA and GAGE were performed using statistically significant differentially expressed genes to determine whether *a priori* defined set of genes were different between the two biological states. For GSEA and GAGE, the molecular Signatures Database v7.3 with the hallmark and canonical (KEGG) gene sets were applied.

### ATAC-Seq and data analysis

Cells from two independent experiments were collected and open chromatin was assessed using an ATACseq kit (Active Motif, 53150). DNAseq library preparation was completed at the Oncogenomics Core Facility at the Sylvester Comprehensive Cancer Center. Samples were sequenced using 100bp paired ends with an Illumina NovaSeq 6000. Compressed Fastq.gz files were uploaded to a Galaxy account (https://usegalaxy.org/), concatenated, and subsequent sequencing reads (∼40 million per sample) were trimmed off the Nextera adapter sequences and filtered using Cutadapt. Reads were mapped to the reference genome (hg38 Canonical) using Bowtie2 with presets “very sensitive end-to-end (--very-sensitive)”, “set the maximum fragment length for valid paired-end alignments: 1000” and allowing mate dovetailing to generate BAM files. We filtered uninformative reads with low mapping quality and were not properly paired using Filter BAM datasets on a variety of attributes (Galaxy Version 2.4.1). Filter on read mapping quality (phred scale) was set to ≥30. ATACseq motif discovery was conducted using HOMER (http://www.homer.ucsd.edu).

### Multi-omics analysis of genes that encode mitochondrial proteins

We examined the differential expression of genes that encode mitochondria-related proteins based on a compendium from MitoCarta (https://www.broadinstitute.org/mitocarta) and mitoXplorer (http://mitoxplorer.ibdm.univ-mrs.fr). We appraised mitochondrial protein-encoding genes using Venn analysis (http://www.interactivenn.net) between the mitochondrial compendium and the differentially regulated genes derived from our RNAseq studies.

### Quantitative real-time RT-PCR (qPCR) and analysis

RNA was prepared like the RNAseq studies. cDNA was synthesized using 200ng total RNA with the ProtoScript® First Strand cDNA Synthesis kit (New England Biolabs) utilizing random hexamers. All cDNAs were amplified under the following conditions: 95°C for 10 minutes to activate AmpliTaq Gold® Polymerase; followed by 40 cycles of 95°C for 15 seconds and 60°C for 1 minute with an internal ROX reference dye. qPCR analysis was performed on a QuantStudio 3 Real-Time instrument (ThermoFisher Scientific) utilizing the Power SYBR™ Green PCR Master mix (ThermoFisher Scientific, **Table S1**). Target genes were normalized to beta actin mRNA expression. For the primer design, the human genome sequence coverage assembly GRCh38.p13 was utilized from the Genome Reference Consortium. Data were presented as fold induction of treatments compared to 0nM (vehicle) normalized to beta actin mRNA levels (i.e., the comparative CT Livak method). Melting curve analysis was performed for all primer sets to eliminate those that yielded primer-dimers. The *p*-values reflect the log fold-change compared to the vehicle (0nM) condition (*n*=3 experimental samples ± SD). A two-way ANOVA test with Sidak’s multiple comparisons test was performed between vehicle and treatment data sets using Prism (GraphPad) where the *p*-value summaries were depicted as ^∗∗∗∗^*p* ≤ 0.0001, ^∗∗∗^*p* ≤ 0.001, ^∗∗^*p* ≤ 0.01, and ^∗^*p* ≤ 0.05.

### MitoSOX Red mitochondrial superoxide indicator and live-cell Apotome imaging

MitoSOX™ Red reagent (ThermoFisher, M36008) was used to detect mitochondrial superoxide levels in live cells. MG-63 cells were cultured in Millicell® EZ chamber slides (EMD Millipore). A 5mM MitoSOX™ Red reagent stock solution was made by dilution into dimethyl sulfoxide (DMSO). A 5μM MitoSOX™ Red reagent working solution was made by diluting the stock into a culture medium. Cells were loaded with MitoSOX™ Red reagent by incubating for 10 minutes at 37°C protected from light. Hoechst 33342 (1:2000) live-cell dye was used as a counterstain to detect the nuclei of live cells (ThermoFisher, H1399). Cells were washed three times with a warm medium. Intensity measurements were obtained using the Zen Blue software (Zeiss) analyzed using Prism 8 (GraphPad). Rotenone (Sigma, R8875) was applied as a positive control. For each replicate (n=6 replicates/condition), average ratios were derived from four different fields of views of 5-10 individual cells. Data are presented as mean±SEM error bars; ^∗∗∗∗^*p* ≤ 0.0001, ^∗∗∗^*p* ≤ 0.001, ^∗∗^*p* ≤ 0.01, and ^∗^*p* ≤ 0.05 (one-way ANOVA with Tukey’s multiple comparisons test compared to vehicle).

### Mitochondrial membrane potential (ΔΨ_M_) measurements and live-cell Apotome imaging

A JC-1 (5,5,6,6’-tetrachloro-1,1’,3,3’ tetraethylbenzimi-dazoylcarbocyanine iodide) mitochondrial membrane potential detection kit (Biotium, 30001) was used to measure mitochondrial membrane potential changes in live cells. MG-63 cells were cultured in Millicell® EZ chamber slides (EMD Millipore). All experiments were performed in a low-light setting. A 1x working solution of JC-1 dye was prepared in a cell culture medium, and cells were incubated in a 37°C cell culture incubator for 15 minutes. Cells were washed once with PBS and replenished with a fresh culture medium. Hoechst 33342 live-cell dye was used to detect the nuclei of live cells (ThermoFisher, H1399). Cells were observed immediately using a Zeiss Observer 7 ApoTome2 microscope using a dual band-pass filter designed to simultaneously detect fluorescein and rhodamine, or fluorescein and Texas Red®. The spectral properties of the JC-1 dye consist of Excitation/Emission (cytoplasm): 510/527 nm (green); and Excitation/Emission (polarized mitochondria): 585/590 nm (red). The ellipsoid spot measurement tool in Imaris was used to select peri-nuclear JC-1-labeled mitochondria to determine JC-1 aggregate:monomer intensity ratios. The spot intensity tool (Imaris) was used to determine the intensity (y-axis) and position (x-axis) between spot-to-spot (i.e., a series of mitochondria-to-mitochondria) in treated cells. For each replicate (n=6-8 replicates/condition), average ratios were derived from four different fields of views of 6-8 individual cells. Data are presented as mean±SEM error bars; ^∗∗∗∗^*p* ≤ 0.0001, ^∗∗∗^*p* ≤ 0.001, ^∗∗^*p* ≤ 0.01, and ^∗^*p* ≤ 0.05 (two-way ANOVA with Sidak’s multiple comparisons test compared to vehicle).

### Mitochondrial biogenesis in-cell ELISA assay

Mitochondrial biogenesis and translation were monitored in MG-63 cells using an in-cell ELISA kit (Abcam, Ab110217). In brief, MG-63 cells were cultured in collagen-coated 96-well plates and treated with vitamin D at various concentrations and durations in five replications. After the treatment series, cells were fixed with 4% PFA and then quenched for endogenous alkaline phosphatase activity using acetic acid. Primary antibody cocktails containing COX-1 and SDHA recognizing antibodies were added to the wells. Afterward, secondary HRP and AP-conjugated antibodies were applied and detected using a microplate reader at OD 405nm (for AP detection of SDHA) and 600nm (for HRP detection of COX-1). Measurements were normalized to the Janus Green staining intensity at OD 595nm to account for differences in cell seeding.

### Immunofluorescence labeling and analysis of MG-63 cells

MG-63 cells were cultured in Millicell® EZ chamber slides (EMD Millipore) and fixed in either 80% methanol or 4% paraformaldehyde (PFA) in 0.1 M phosphate buffer (PBS, pH 7.4) for 10 minutes. PFA fixed cells were permeabilized with 0.2% Triton X-100 in PBS for five-to-fifteen min at room temperature, followed by washes with PBS. Cells were blocked with normal horse/goat serum for non-specific background, and then incubated with primary antibodies at a 1:200 dilution for one hour at room temperature. Primary antibodies used in this study included: Rabbit monoclonal to VDAC1 (Abcam, ab154856), and rabbit monoclonal to REDD1/DDIT4 (Abcam, ab191871). Following washing steps in phosphate-buffered saline-Tween-20, the cells were incubated at room temperature for 20 min with corresponding species-specific secondary antibodies (Alexa series at 1:2000, Life Technologies). The slides were covered with Vectashield medium containing 4′,6-diamidino-2-phenylindole (DAPI; Vector Laboratories, H- 1200-10) for nuclei staining and then mounted with a glass coverslip. Negative controls were included that had either no primary or secondary antibodies in the blocking buffer.

Immunofluorescence confocal-like microscopy was performed using a Zeiss Observer 7 ApoTome2 system. We carefully selected the wavelength ranges, emission filters, and dichroic mirrors to avoid signal bleed-through. Deconvolution and Apotome processing (i.e., extended depth of view) of stacked images was performed using the ZenBlue software (Zeiss Microscope). Image stacks were reconstructed and visualized as three-dimensional (3D) volumes with Imaris software (Bitplane). The Imaris Spot detection algorithm was used as described by the manufacturer for semiautomatic identification and counting of fluorescently labeled mitochondria and cytoplasmic components. Means of expression intensity of mitochondrial and cytoplasmic regions within individual cells were compared between vehicle and vitamin D-treated samples. For analysis, individual experiments (n=4) were performed whereby each experiment entailed an assessment of 4-6 individual sets of cells for technical replication. For some experiments, Imaris (Bitplane) and MATLAB were used to generate 3D rendered models of protein expression and co-localization. Spots are located at the local maxima of the filtered image with background subtraction. Imaris calculated a ‘spot quality’ (minimum of 100) based on intensity differences and shapes for spot rendering and was adjusted to include the signal of interest. Co-localization analysis was performed using the “spot” tool to designate the distance threshold and the mean distance between the “VDAC” and “DDIT4” co-localized spots. A two-way ANOVA test with Sidak’s multiple comparisons test was performed between vehicle and treatment data sets using Prism (GraphPad) where the *p*-value summaries were depicted as ^∗∗∗∗^*p* ≤ 0.0001, ^∗∗∗^*p* ≤ 0.001, ^∗∗^*p* ≤ 0.01, and ^∗^*p* ≤ 0.05. Statistical significance was accepted at ^∗^*p* ≤ 0.05.

### Transmission electron microscopy (TEM)

TEM was performed at the Transmission Electron Microscopy Core Facility at the Miller School of Medicine, University of Miami. The TEM Core prepared the cells for electron microscopy and performed embedding and semi-thin (1µm) and thin (100nm) sectioning of the samples and final imaging with a JEOL JEM-1400 electron microscope. For analysis, between 7-10 cells were investigated per condition, in which we averaged parameters between 20-40 mitochondria per cell.

### Stimulation and measurement of ER stress

Known ER stress inducers tunicamycin (Sigma-Aldrich, T7765) and thapsigargin (Sigma-Aldrich, T9033) were diluted in ethanol and exposed to cells for 6 hours with appropriate vehicle controls. Both end-point semi-quantitative and quantitative real-time PCR methods were used to assess ER stress base on Yoon Seung-Bin et al. 2019 with adjustments (e.g., the annealing temperature of 62°C was used instead). For the end-point PCR reaction, the Phusion DNA polymerase (ThermoFisher) was used, and a 2.5% agarose gel was utilized to assess ER stress PCR products. u/s/tXBP1 primers relative to the 26bp of XBP1 removed by IRE1 were used for real-time PCR reactions (**Table S1**). Housekeeping genes (Gapdh, 18sRNA) and the total amount of XBP1 (**Table S1**) were used to normalize gene expression.

## Supplemental Figures

**Figure S1.**
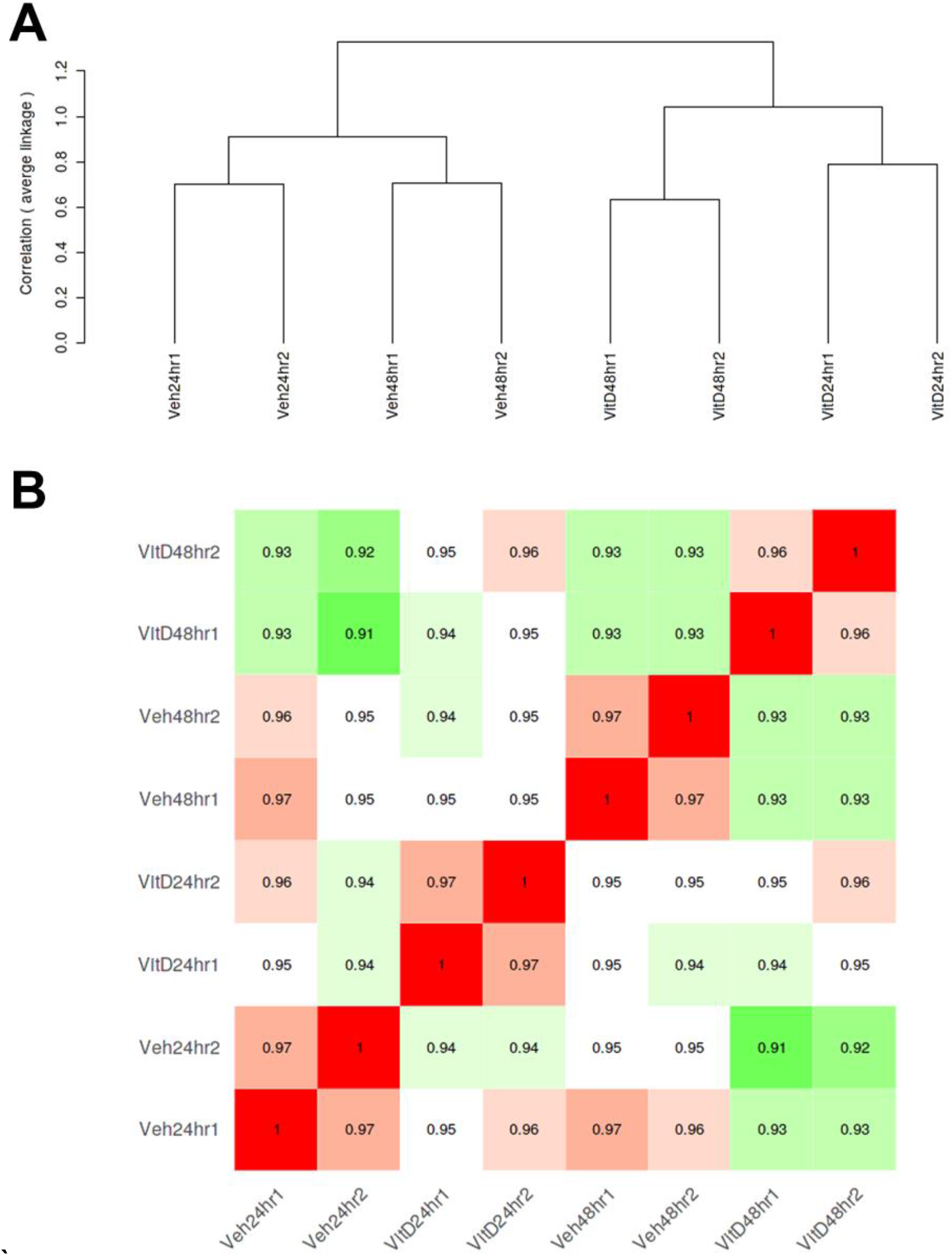
**Correlation matrix of top 75% of RNAseq transcripts** A) Hierarchical clustering tree. The tree generated using genes with maximum expression level at the top 75%. B) Pearson’s correlation coefficients across all data sets.

**Figure S2.**
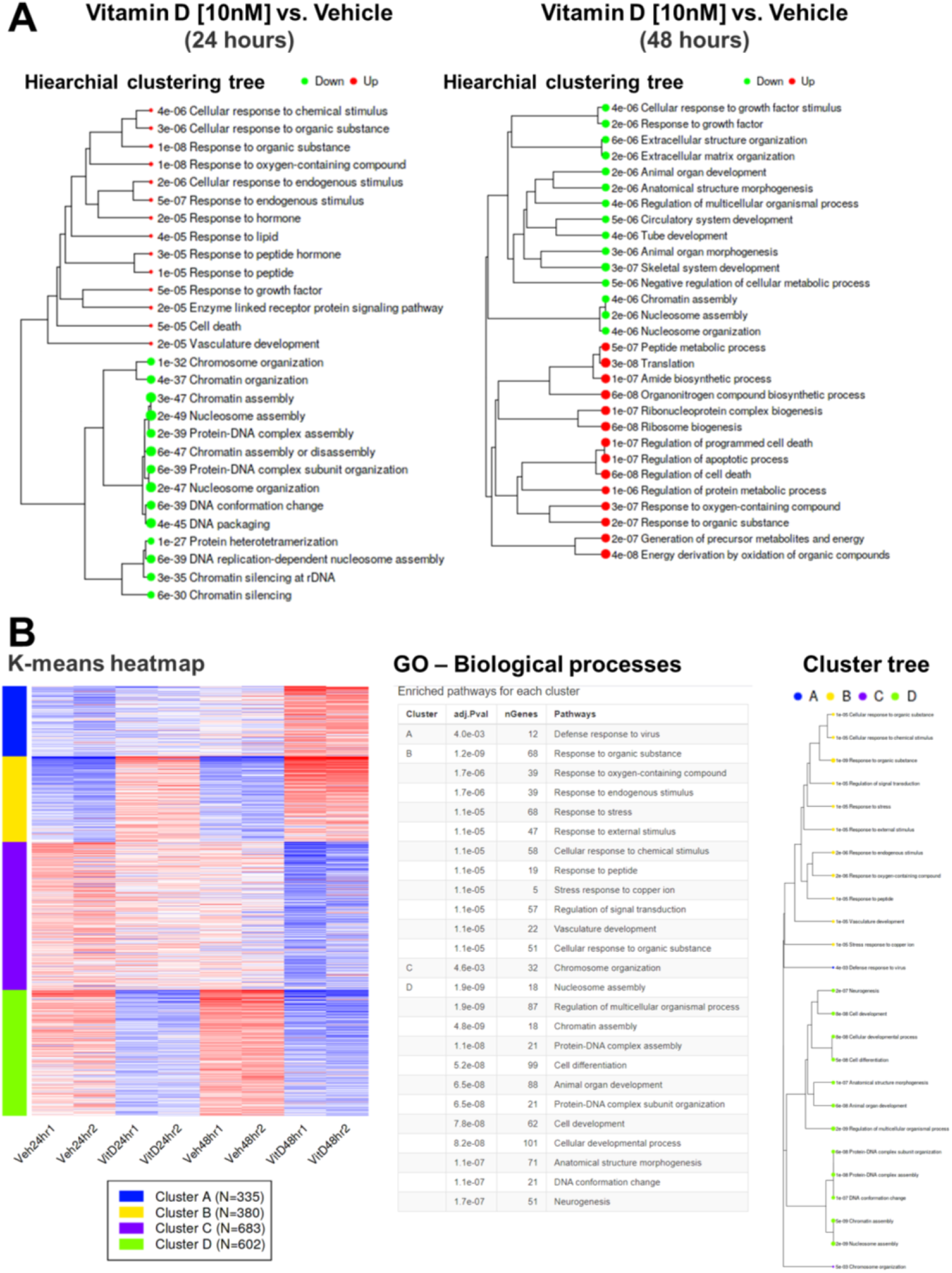
**Hierarchical and K-means clustering of RNAseq data sets** A) Visualization of the relationships/correlations among enriched GO terms using hierarchical clustering tree using iDEP. For the tree construction, we first measured the distance among the GO terms by the percentage of overlapped genes. B) For K-means clustering, we used the dimension reduction algorithm t-SNE to map the top 2000 most variable genes, and then examined the distribution to help choose the number of clusters in K-means. For heatmap generation, we normalized by gene mean center and then mapped back to the GO terms to further generate tables and cluster tree.

**Figure S3.**
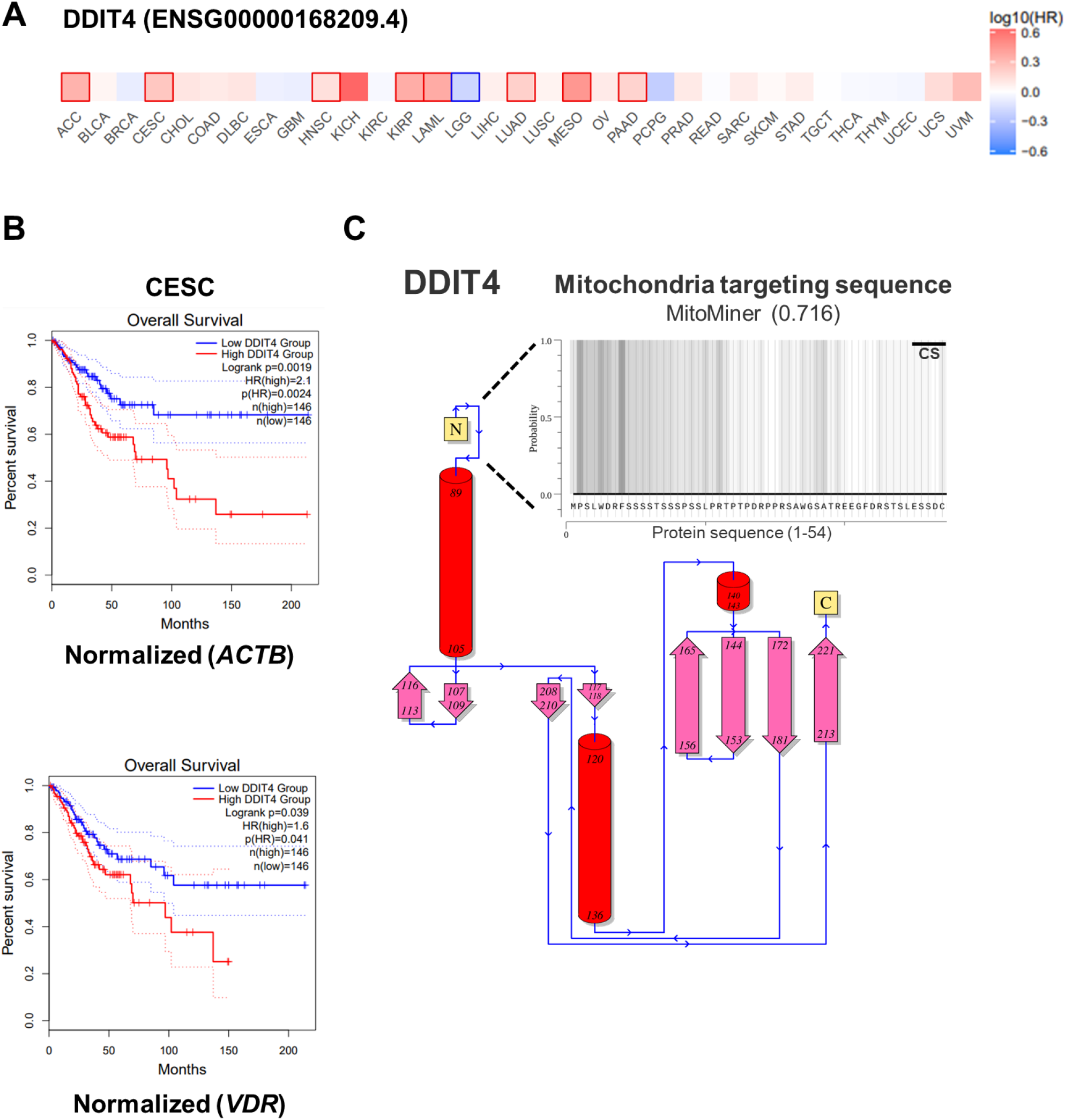
**DDIT4 in cancer and mitochondria** A) Gene Expression Profiling Interactive Analysis (GEPIA), a meta-analysis of individual cancer data sets, shows that DDIT4 mRNA expression is significantly increased in numerous tumor tissues such as adrenocortical carcinoma, cervical squamous cell carcinoma (CESC), head and neck squamous cell carcinoma, kidney renal papillary cell carcinoma, acute myeloid leukemia, lung adenocarcinoma, mesothelioma, and pancreatic adenocarcinoma. No data on osteosarcoma is available through GEPIA. B) GEPIA was used to determine the overall cancer survival for CESC based on *DDIT4* gene expression levels. DDIT4 levels were normalized for relative comparison between a housekeeping gene (*ACTB*) and the *VDR* gene. Using the log-rank test (Mantel-Cox test) for hypothesis evaluation, the hazard ratio (HR) and the 95% confidence interval information were included in the survival plots for the high and low expressing cohorts. C) *In silico* approach to identify putative mitochondria targeting sequences in the proximal region of the human DDIT4 protein (UniProt: Q9NX09) using MitoMiner (https://mitominer.mrc-mbu.cam.ac.uk/release-4.0). Based on the amino acid sequence of DDIT4, MitoMiner predicted a mitochondrial targeting sequence with an overall probability score of 0.716 along with the prediction of a cleavage sequence in the N-terminal region.

## Supplemental Table

**Supplemental Table S1.**
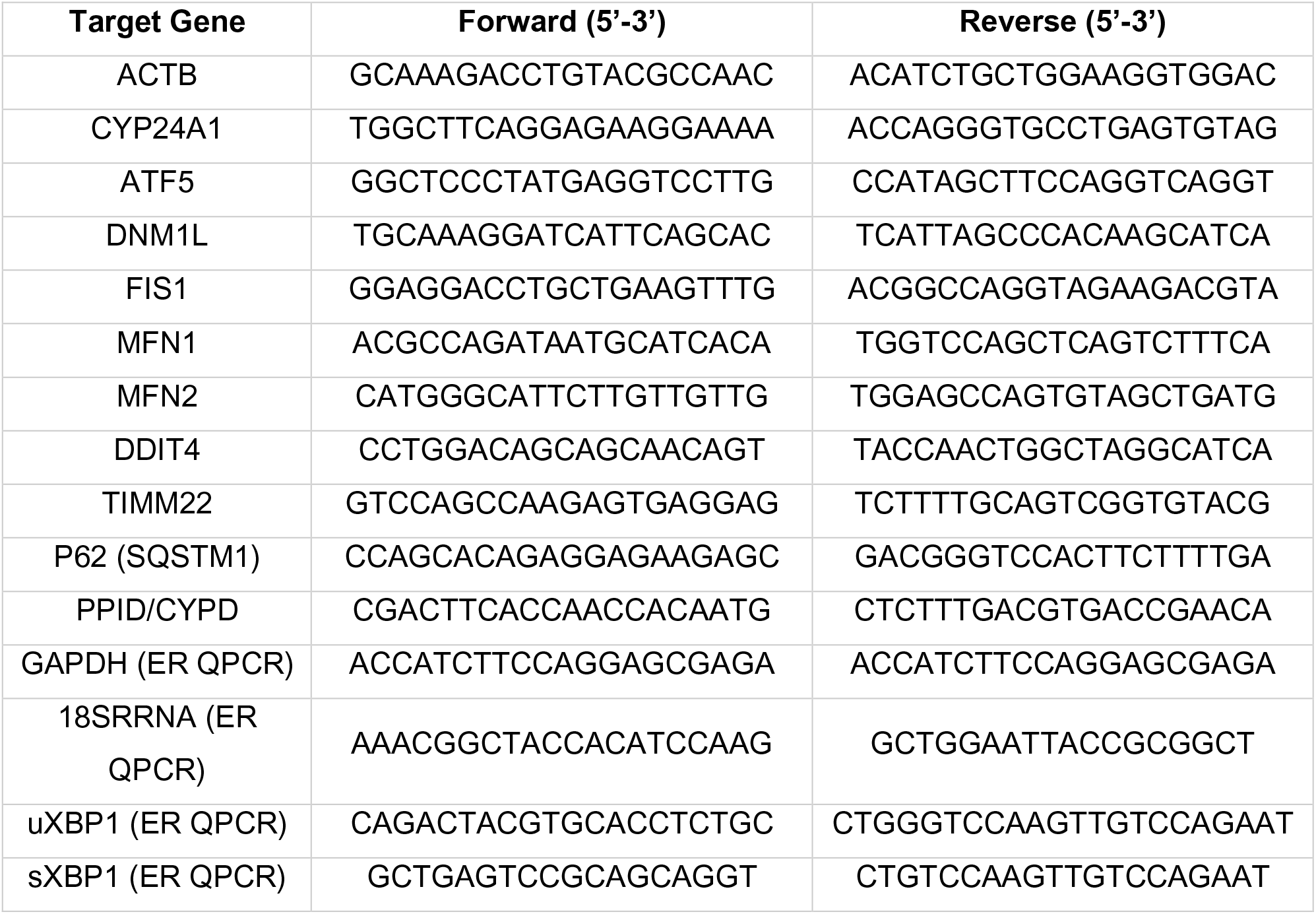
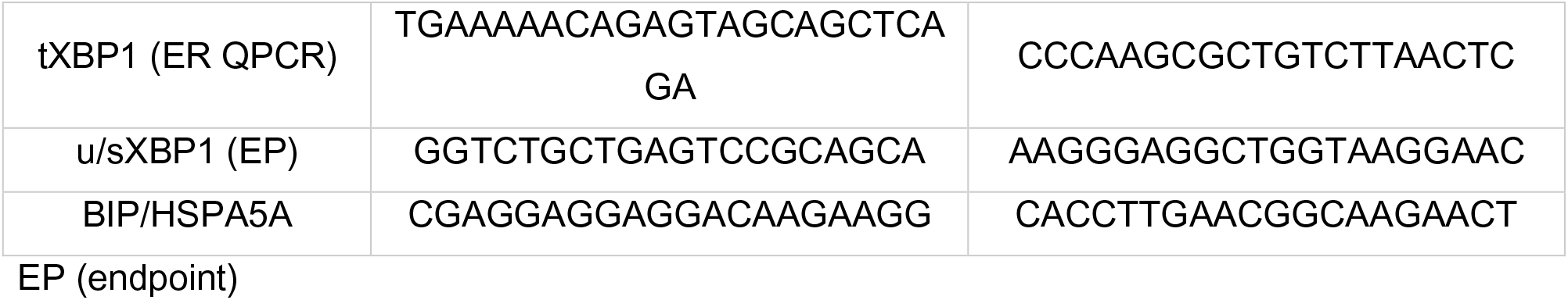
. Human real-time PCR and end-point primer sets

